# Novel functional sequences uncovered through a bovine multi-assembly graph

**DOI:** 10.1101/2021.01.08.425845

**Authors:** Danang Crysnanto, Alexander S. Leonard, Zih-Hua Fang, Hubert Pausch

## Abstract

Linear reference genomes are typically assembled from single individuals. They are unable to reflect the genetic diversity of populations and lack millions of bases. To overcome such limitations and make non-reference sequences amenable to genetic investigations, we build a multi-assembly graph from six reference-quality assemblies from taurine cattle and their close relatives. We uncover 70,329,827 bases that are missing in the bovine linear reference genome. The missing sequences encode novel transcripts that are differentially expressed between individual animals. Reads which were previously poorly or unmapped against the bovine reference genome now align accurately to the non-reference sequences. We show that the non-reference sequences contain polymorphic sites that segregate within and between breeds of cattle. Our efforts to uncover novel functional sequences from a multi-assembly graph pave the way towards the transition to a more representative bovine reference genome.

## Introduction

A well-annotated reference genome enables systematic characterization of sequence variation within and between populations, as well as across species. The reference genome of domestic cattle (*Bos taurus taurus*) was generated from the inbred Hereford cow «L1 Dominette 01449»^1^. Long-read sequencing and sophisticated genome assembly methods have enabled spectacular improvements in the contiguity and quality of the bovine reference genome. The contig (contiguous sequence formed by overlapping reads without gaps) N50 size (i.e., 50% of the genome is in contigs of this size or greater) of the bovine reference genome has increased from kilo- to megabases over the past five years^2^. Recent method and sequencing technology developments have facilitated the assembly of multiple reference-quality genomes. The application of trio-binning^3^ resulted in chromosome-scale haplotype-resolved assemblies for three taurine (Hereford, Angus, Highland cattle) and one indicine (Brahman) cattle breeds, as well as for yak (*Bos grunniens*), a closely related species to domestic cattle^4,5^.

Bovine DNA sequences are typically aligned to the Hereford-based reference genome to discover and genotype variable sites. Reference-guided read alignment and variant genotyping has revealed millions of polymorphic variants that segregate within and between taurine and indicine cattle breeds^6–8^. However, using the linear reference in this alignment approach is susceptible to reference allele bias, particularly for DNA samples that are greatly diverged from the reference^9,10^. Moreover, reference-guided methods are blind to variations in sequences that are not present in the reference genome^11^. Recent estimates suggest that millions of bases are missing in mammalian reference genomes^12,13^, indicating a high potential for bias.

Efforts to mitigate reference allele bias and increase the genetic diversity of reference genomes have led to graph-based references^14,15^. We have previously shown that a bovine genome graph, which integrates linear reference coordinates and pre-selected variants, improves the mapping of reads and enables unbiased variant genotyping^16,17^. However, previous attempts focused on augmenting the bovine reference genome with small variations (<50bp), not the larger class of structural variations. Despite being an important source of genotypic and phenotypic diversity^18,19^, little is known about the prevalence and functional impact of structural variations in the cattle genome. The availability of reference-quality assemblies and long read sequencing data from different breeds of cattle now provides an opportunity to characterize sequence diversity beyond small variations^20,21^.

In this paper, we integrate reference-quality assemblies from multiple taurine breeds as well as two close relatives into the first bovine multi-assembly graph with minigraph^21^. We detect autosomal sequences that are missing in the bovine reference genome and investigate their functional significance using transcriptome data. We show that the non-reference sequences contain novel transcripts that are differentially expressed as well as polymorphic sites that segregate within and between breeds of cattle.

## Results

### Construction of a bovine multi-assembly graph

We considered the Hereford-based bovine reference genome and five reference-quality assemblies from three breeds of taurine (*Bos taurus taurus*) cattle (Angus, Highland, Original Braunvieh)^2,4,5^ and their close relatives Brahman (*Bos taurus indicus*)^4^ and yak (*Bos grunniens*)^5^. All assemblies, except for the Original Braunvieh, were generated prior to this study. The reference-quality assembly for an Original Braunvieh female calf was created with 28-fold PacBio HiFi read coverage (see Supplementary Note 1). The contig and scaffold N50 values of the six assemblies ranged from 21 to 80 Mb and 86.2 to 108 Mb, respectively (Table 1).

**Table 1:** Details of six bovine genome assemblies

| Breed | Sex<sup>1</sup> | Primary data used for the assembly<sup>2</sup> | Type of assembly | Assembler | Contig N50 (Mb) | Scaffold N50 (Mb) | Length of the autosomes |
| --- | --- | --- | --- | --- | --- | --- | --- |
| Hereford – *Bos taurus taurus* | F | PacBio (80-fold CLR) | Primary | Falcon | 21 | 108 | 2,489,385,779 |
| Angus - *Bos taurus taurus* | M | PacBio (136-fold CLR) | Haplotype-resolved | TrioCanu | 29.4 | 102.8 | 2,468,157,877 |
| Highland - *Bos taurus taurus* | F | PacBio (125-fold CLR) | Haplotype-resolved | TrioCanu | 71.7 | 86.2 | 2,483,452,092 |
| Original Braunvieh - *Bos taurus taurus* | F | PacBio (28-fold HiFi) | Primary | Hifiasm | 86.0 | 96.3 | 2,607,746,442 |
| Brahman - *Bos taurus indicus* | F | PacBio (136-fold CLR) | Haplotype-resolved | TrioCanu | 23.4 | 104.5 | 2,478,073,158 |
| Yak – *Bos grunniens* | F | PacBio (125-fold CLR) | Haplotype-resolved | TrioCanu | 70.9 | 94.7 | 2,478,308,164 |

<sup>1</sup> Female (F) and male (M) assemblies contain either X or Y chromosomal sequences
<sup>2</sup> Additional data may have been used to polish the assemblies and facilitate scaffolding; CLR: continuous long reads; HiFi: high-fidelity

The six assemblies were integrated into a multi-assembly graph with minigraph. We only considered autosomal sequences because the haplotype-resolved assemblies represent either paternal or maternal haplotypes, thus lacking either X or Y chromosomal sequences. The Hereford-based linear reference genome (ARS-UCD1.2) formed the backbone of the bovine multi-assembly graph. The graph was then augmented with the five additional assemblies, added in order of increasing Mash-distance from the ARS-UCD1.2 reference^22^ (Fig. 1). Constructing this multi-assembly graph took 4.1 CPU hours and 58 GB of RAM, taking 36 minutes of wall-clock time when using 10 threads.

**Fig. 1:**
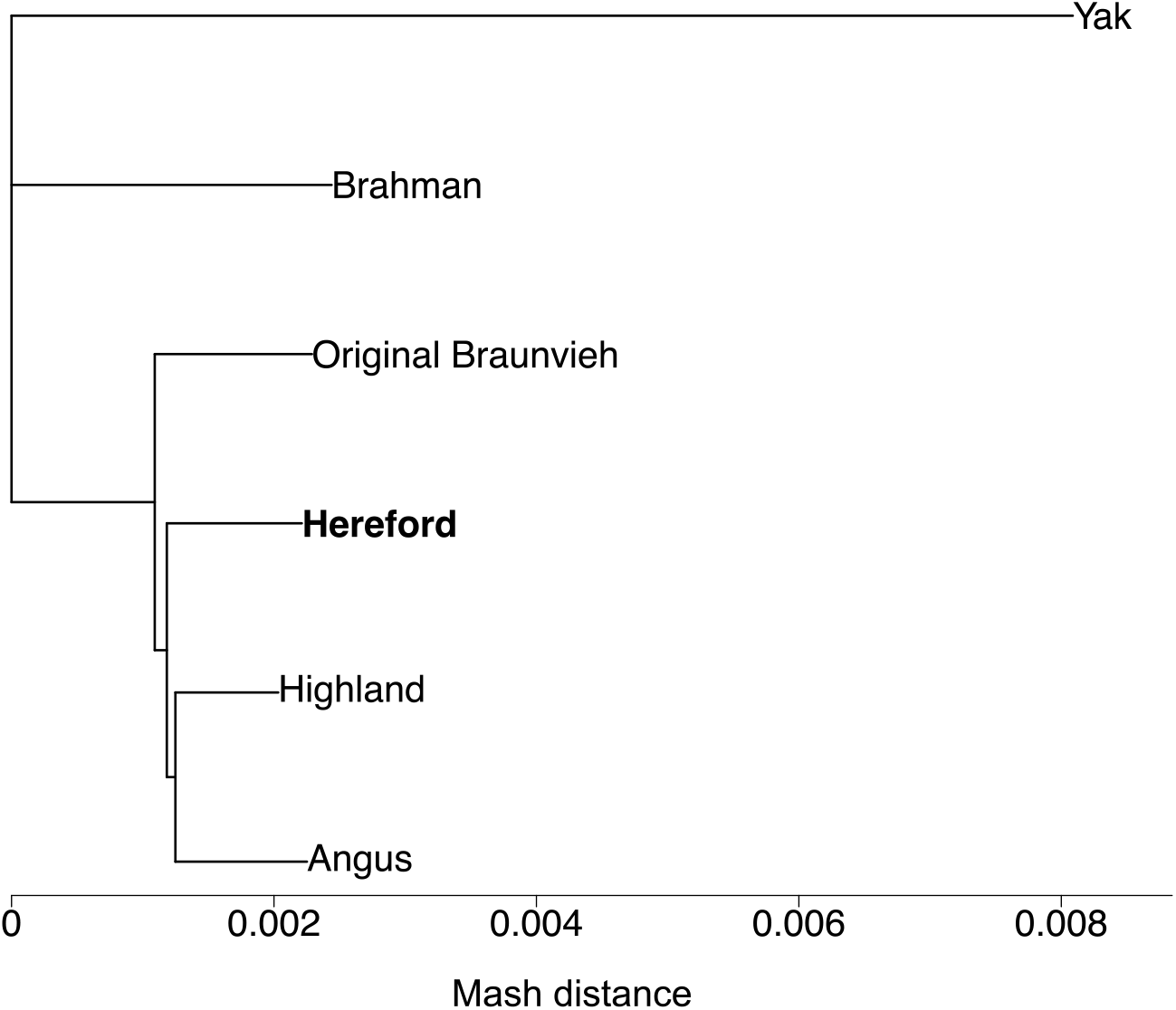
Phylogenetic distance between six genome assemblies. A Mash-based phylogenetic tree derived from six bovine assemblies, including the current Hereford-based reference genome (bold). The yak assembly was used as the outgroup to root the tree during building.

### Recovery of non-reference sequences from the multi-assembly graph

Our bovine multi-assembly graph represents 2,558,596,439 nucleotides, spread across 182,940 nodes connected by 258,396 edges. On average, a node spans 13,985 nucleotides and is connected by 1.4 edges. Of the edges, 141,086, 113,332, and 3,978 connect two reference nodes, a reference and non-reference node, or two non-reference nodes, respectively.

The vast majority (2,489,385,779 or 97.29%) of nucleotides in the multi-assembly graph originate from the linear reference backbone, covered in 123,483 nodes. These reference nodes span 23,088 bases on average, ranging from 100 to 1,398,882 bases. The incremental integration of the Highland, Angus, Original Braunvieh, Brahman, and yak assemblies added 8,847, 4,613, 3,555, 11,996, and 30,446 non-reference nodes, respectively containing 14,679,286, 5,537,769, 7,013,258, 11,116,220, and 30,864,127 non-reference bases. The resulting multi-assembly graph contained 59,457 non-reference nodes spanning 69,210,660 bases.

To determine the support of the non-reference nodes, we aligned individual assemblies back to the multi-assembly graph. Nodes were then labelled according to which assembly path traversed them (see Supplementary Figs. 1 & 2). This approach also enabled a straightforward confirmation of minigraph’s mapping accuracy. Only reference nodes should contain a Hereford label, since this assembly was used as the backbone of the graph. Mapping was highly accurate, as indicated by an F1 score of 99.97%.

The non-reference nodes had a cumulative length of 43,341,418, 23,644,772, 18,202,102, 14,453,112 and 15,542,368 bases in the yak, Brahman, Original Braunvieh, Angus, and Highland assemblies. Most non-reference nodes (41,855 or 70.40%) and non-reference sequences (42.52 Mb, 69.52%) were either private to yak (29,854 nodes, 29.9 Mb), Brahman (7,843 nodes, 8.22 Mb), or shared by both assemblies (4,158 nodes, 3.05 Mb) (Fig. 2). The Original Braunvieh, Highland, and Angus assemblies contributed 4.51, 2.78 and 2.39 Mb in 2,016, 1,938 and 1,759 nodes, respectively, that were not detected in any other assembly. The three taurine assemblies shared 668 nodes containing 0.77 Mb not detected in ARS-UCD1.2, yak, or Brahman. There were also 1,318 non-reference nodes with a cumulative length of 4.4 Mb supported by all five additional assemblies.

**Fig. 2:**
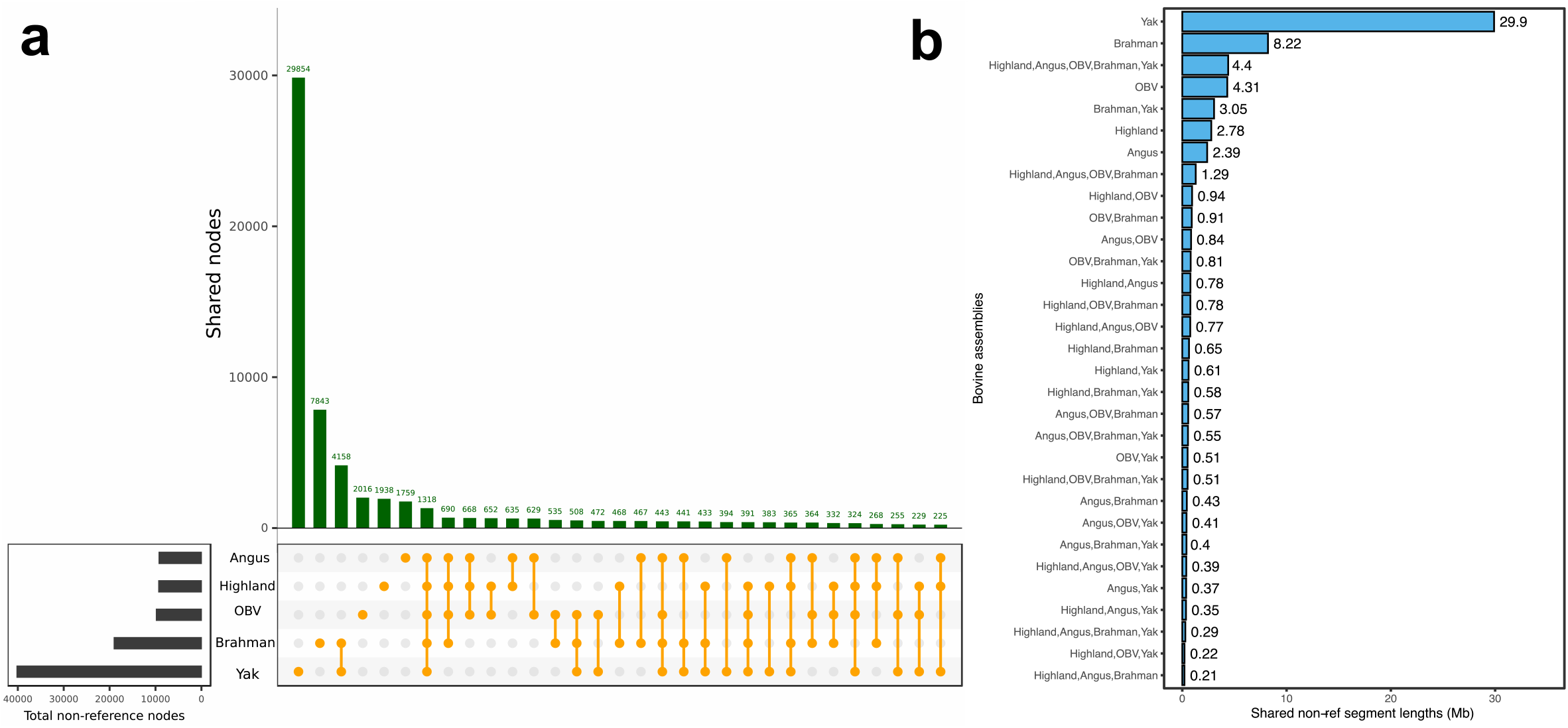
Non-reference sequences detected across assemblies. Intersection of non-reference nodes (a) and cumulative length of non-reference sequences (b) found in five assemblies. OBV = Original Braunvieh.

The core genome of the multi-assembly graph (i.e., nodes shared by all assemblies) is contained in 67,482 nodes with a cumulative length of 2,402,561,410 bases. About 6.10% of the pangenome (115,458 nodes containing 156,035,029 bases) is flexible (i.e., not shared by all assemblies). Of the flexible part, 69,697 nodes containing 97,106,100 bases are shared by at least two assemblies, and 45,761 nodes with 58,928,929 bases are only found in one assembly. The profile of the multi-assembly graph changes markedly when distant assemblies (e.g., Brahman, yak) are added (Supplementary Note 2).

### Structural variation discovery from the multi-assembly graph

Using the bubble popping algorithm of gfatools^21^, we identified 68,328 structural variations present in the multi-assembly graph. To reveal true alleles within these structural variations, we traversed all possible paths through the bubbles (i.e., alleles) and retained only those that were supported by at least one assembly (Supplementary Fig. 2). Most of the structural variations had two alleles (64,224 or 94%). The remaining 4,104 structural variations were multi-allelic, most of which had three alleles (3,324 or 81%). We identified 141,747 alleles at the structural variations, including 73,506 non-reference alleles with a cumulative length of 74,453,929 bases.

We overlapped the breakpoints of the structural variations with the Ensembl annotation (build 101) of the bovine reference genome. Almost all structural variations are either intergenic (47,642 or 69.81%) or intronic (20,227 or 29.64%). There were 338 unique genes affected by structural variations, with 170 and 202 disrupting coding sequences and exons respectively. A Panther GO-Slim Biological Process^23^ analysis indicated that these genes are enriched for genes related to the adaptive immune response (4.35-fold, *P*=0.04), T-cell mediated immunity (6.37-fold, *P*=0.04), actin filament depolymerization (8.54-fold, *P*=6.56e-03), microtubule cytoskeleton organization (10.48-fold, *P*=1.85e-04), and iron sulfur cluster assembly (9.96-fold, *P*=0.02).

The non-reference alleles consisted of 40,369 insertions and 33,137 deletions with an average length of 1,181 and 1,210 bases respectively (Supplementary Table 1). The cumulative length (absolute difference between reference and non-reference allele) was longer for insertions (47,691,942 bases) than deletions (40,101,303 bases). This pattern was similar for biallelic variations (35,748 and 28,476 biallelic insertions and deletions, respectively, encompassing 37,388,222 and 28,373,582 bases with an average variant length of 1,045 and 996 bases). The multi-assembly graph contained more complete insertions (20,432; i.e., only non-reference sequences present in the bubbles, thus reference length is 0) than alternate insertions (15,316; i.e., both reference and non-reference sequences present but non-reference allele is longer). The pattern was similar for deletions. The multi-allelic structural variations had 13,299 alleles including 9,282 non-reference alleles with 4,621 insertions and 4,661 deletions, respectively, affecting 11,727,721 and 10,303,720 bases. Bubbles with multi-allelic structural variations contained more mixed mutations (1,941; both deletions and insertions detected within the same bubble) than multiple mutations of the same type (994 and 1,082 for multiple insertions and deletions, respectively).

The yak, Brahman, Original Braunvieh, Angus, and Highland assembly contained respectively 49,836, 22,976, 10,965, 10,735, and 10,560 non-reference alleles (Supplementary Fig. 3). Most non-reference alleles (36,443, total length: 30 Mb) were private to the yak assembly. We detected 9,267, 2,232, 2,133, and 2,037 non-reference alleles, respectively, containing 10.2, 4.9, 3.8, and 3.3 Mb that were private to the Brahman, Original Braunvieh, Highland, and Angus assembly. All assemblies contained more insertions than deletions relative to the ARS-UCD1.2 reference (e.g., 5,537 (8.57 Mb) insertion alleles versus 3,730 (1.5 Mb) deletion alleles were private to the Brahman assembly). We also found 1,749 alleles within the 4.4 Mb of non-reference sequence (2.1 Mb of which is non-repetitive) shared by all assemblies except ARS-UCD1.2.

### Validation of the structural variations using long sequencing reads

We mapped 6,803,270 PacBio HiFi reads with a mean length of 20,612 bases (~46-fold coverage) from a Nelore (*Bos taurus indicus*) x Brown Swiss (*Bos taurus taurus*) crossbred bull to the multi-assembly graph using GraphAligner^24^. Neither parental breed is present in the multi-assembly graph.

To assess the support for the structural variations derived from the multi-assembly graph, we calculated the HiFi read coverage (measured as the number of reads aligned) at each node and edge in the multi-assembly graph. Of 59,457 non-reference nodes, 23,418 (39.38%) were strongly supported, being covered by more than 10 reads. The aligned HiFi reads traversed 37,288 edges that connect 23,418 non-reference and reference nodes, implying that 32.90% of 113,332 structural variation breakpoints of the multi-assembly graph had support from the hybrid cattle.

We then investigated the relative support from the HiFi reads for the reference and non-reference alleles in the 68,328 bubbles. Nearly half (35,247 or 48.0%) of the non-reference alleles were supported by the crossbred bull. Of these, 8,647 were more supported than the corresponding reference allele. In 3,849 of these cases, only the non-reference allele was supported, with no reads aligning to the reference allele.

### Sequence content of the structural variations

In order to investigate the functional relevance of the non-reference sequences, we extracted 45,357 non-reference alleles from the 70,329,827 non-reference bases in the multi-assembly graph (Supplementary Fig. 4). These sequences originate from 38,906 biallelic and 6,451 multiallelic structural variations, respectively, that have a cumulative length of 43,003,591 and 27,326,236 bases. On average, the alleles of multiallelic structural variations were four times longer than that of biallelic bubbles (4,205 versus 1,104 bases).

The non-reference sequences are largely comprised of repetitive elements (53,690,260 bases or 76.34%, Supplementary Fig. 5). LINE/L1 and LINE/RTE-BovB account for 28.04 (52.22%) and 6.77 (12.61%) Mb of repetitive non-reference sequence, respectively.

Repetitive sequences (both interspersed and simple repeats) are more evenly distributed across the autosomes than non-repetitive sequences. Many repetitive and non-repetitive non-reference sequences were detected at two regions on bovine chromosomes 18 and 23 that encompass the leukocyte receptor complex and the major histocompatibility complex (Supplementary Fig. 6).

We hypothesized that the 16,639,567 non-repetitive non-reference bases contain transcribed sequences. A BLASTX search of these sequences against a protein sequence database of *Bos* and related species revealed hits for 403 structural variations containing 299,337 non-reference bases. As a complementary approach, we predicted genes from the non-repetitive sequences using the Augustus software tool. The *ab initio* prediction revealed 857 gene models from 768 distinct structural variations that had a minimum coding sequence length of 150 bp, including 374 complete gene models with transcription start site, start codon, exons, stop codon, and transcription termination site (Supplementary Table 2). On average, the transcript, coding sequence, and protein length of the complete gene models is respectively 4,742 bp, 794 bp, and 264 aa.

### *De novo* transcript assembly from the non-reference sequences

As the two complementary gene prediction methods indicated that the novel sequences contain transcribed features, we sought experimental evidence. We appended the 70 Mb of repeat masked non-reference sequences contained in 45,357 additional contigs to the ARS-UCD1.2 reference, making an extended bovine reference genome. This renders the non-reference sequences amenable to current methods of linear mapping of transcriptome data. Using HISAT2, we aligned liver transcriptomes from 39 cattle across Angus, Holstein, Jersey, and Brahman breeds to both the linear reference as well as the extended reference. We also aligned transcriptomes from Dominette, the animal sequenced to assemble the reference genome. A greater portion of reads mapped to the extended reference compared to the original reference for all examined samples (Supplementary Fig. 7). Across the 40 samples, the overall mapping rate increased by 0.037%, which corresponds to approximately 18K reads for a paired-end RNA-seq dataset of 25 million reads. The mapping improvements were larger for samples with great genetic distance from the reference genome. Brahman had the largest improvement (0.060%), followed by the taurine breeds: Angus (0.032%), Holstein (0.026%), and Jersey (0.030%). As expected, Dominette benefitted the least (0.010%), but still demonstrated an improvement over using the original reference.

Next, we used StringTie2^25^, guided with gene models predicted by Augustus (see above), to assemble reads which aligned to non-reference sequences into 1,431 putatively novel genes. Of these, 885 were expressed at TPM ≥ 1 in at least one breed, including 405 that were originally predicted by Augustus. We selected these 405 putatively novel genes, supported by both *ab initio* prediction and *de novo* transcript assembly for further analyses.

A subset of 263 putatively novel genes were also expressed at TPM ≥ 1 in liver transcriptome data from Dominette. Many of these genes might be divergent copies of ribosomal proteins and olfactory receptors, as indicated by BLASTP queries. Expression of 142 putatively novel genes was detected between 1 and 1574 (mean=21) TPM in liver transcriptomes of Angus, Holstein, Jersey, or Brahman cattle, but not in Dominette. These were mostly expressed in Brahman cattle (75 genes per animal, 117 across all Brahman animals, Fig. 3a) including 20 that were specific to this breed. Among the taurine breeds, we detected more novel genes at ≥1 TPM in Angus (N=109) than either Holstein or Jersey (both N=84) cattle. Putatively novel genes common to all four non-reference breeds accounted for nearly half, 68 of the 142, identified in any non-reference breed (Fig. 3b). The average expression was significantly higher (*P*=0.004, one-tailed t-test) for genes that were expressed in at least two breeds (N=106, TPM=13.48) than genes expressed in only one breed (N=36, TPM=1.64). BLASTP queries provided additional support for 57 out of 142 putatively novel genes (Supplementary Fig. 8). The top hits suggest that the putatively novel genes encode proteins related to: immune response (antigen-presenting glycoprotein, immunoglobulin, BOLA, killer-T-cell, interferon, Ig-like lectin, CMRF35, MHC, cytokine), signalling (G-protein signalling protein, tyrosine-phosphatase), cytoskeleton regulations (myosin, actin, twinfilin, KANTB1), lipid metabolism (apolipoprotein, lipid-binding protein), and protein modifications (heat-shock chaperone, ubiquitin conjugating enzyme, rhoA ubiquitin).

**Figure 3.**
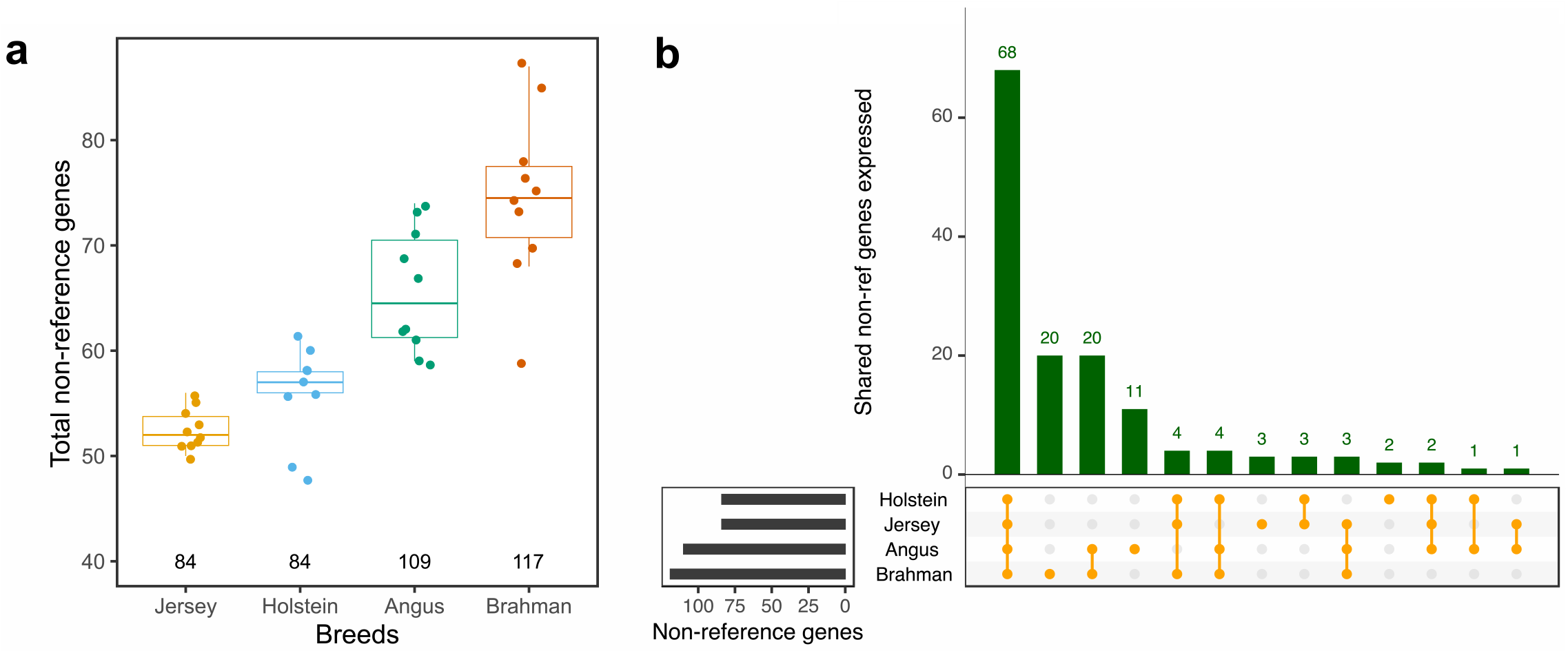
Novel transcribed genes detected from non-reference sequences. (a) Number of non-reference genes expressed ≥1 TPM in liver tissue from taurine (Jersey, Holstein, Angus) and indicine (Brahman) cattle breeds. Each point represents the number of novel genes detected per animal. The number of distinct novel genes detected for each breed is indicated below the boxplots. (b) Expression of 142 putatively novel non-reference genes in the four cattle breeds.

### Non-reference sequences contain differentially expressed genes

To investigate if the non-repetitive sequences also encode transcripts that are differentially expressed between individual animals, we obtained publicly available peripheral blood leukocyte transcriptome data for eight *Mycobacterium bovis*-infected and eight non-infected Holstein cattle^26^. Following the transcriptome analysis introduced earlier, between 9,255,069 and 25,310,808 RNA sequencing reads aligned to the extended bovine reference genome. The subsequent *de novo* transcript assembly produced 949 transcripts, encoded by 661 putatively novel genes. We appended them to the Ensembl ARS-UCD1.2 reference genome annotation, yielding a total of 28,268 genes. We detected expression levels ≥ 1 CPM in at least eight samples for 13,085 genes, including 272 novel genes. We subsequently tested these genes for differential expression, finding 3,646 genes, including 36 putatively novel genes, which were differentially expressed (FDR ≤ 0.05) between *Mycobacterium bovis*-infected and non-infected cattle (Fig. 4a). The top differentially expressed genes from the ARS-UCD1.2 annotation, as well as their transcript abundances in cases and controls, agreed well with the original findings from McLoughlin *et al.*^26^ that were based on the previous UMD3.1 annotation (Supplementary Note 3).

**Fig. 4.**
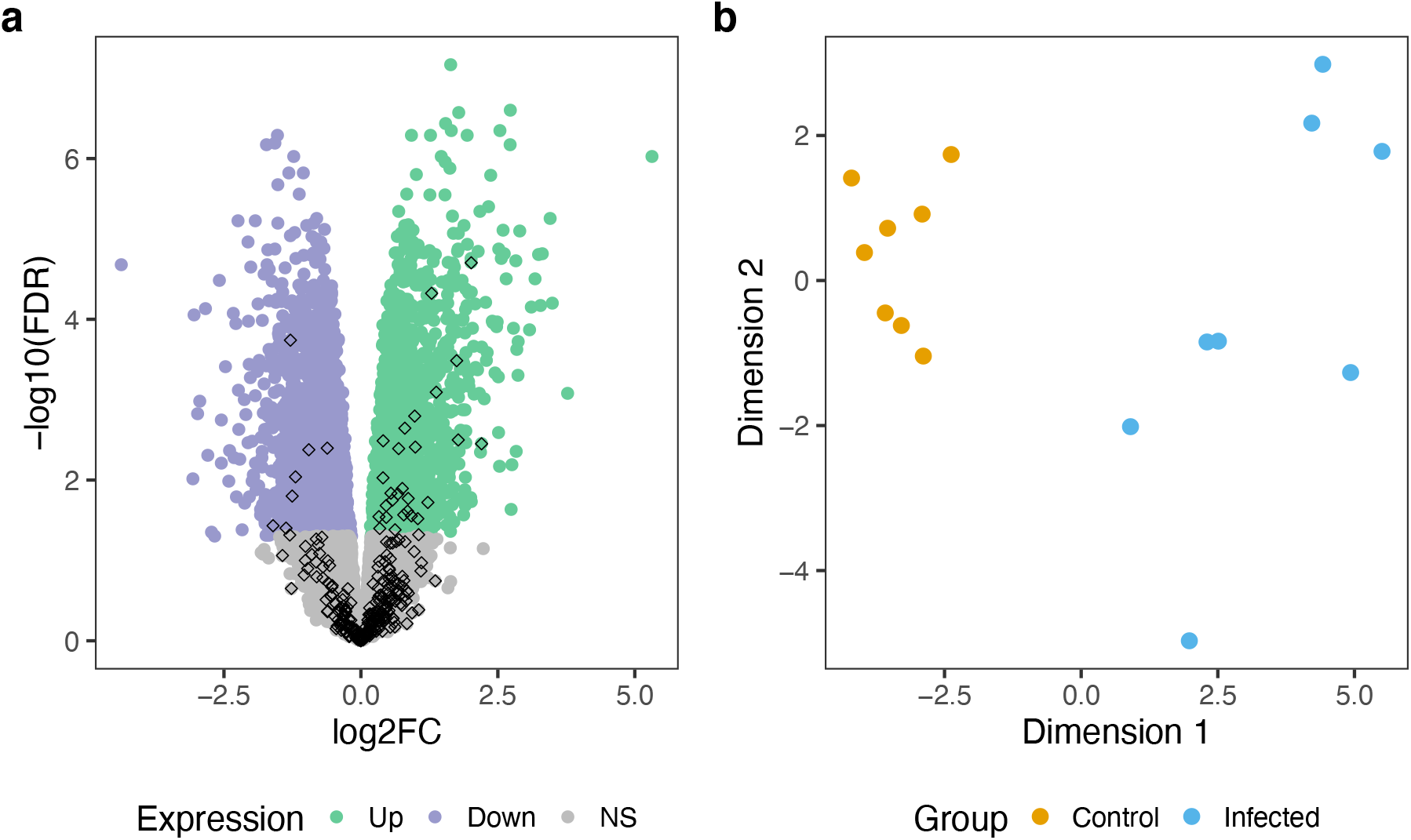
Differentially expressed non-reference genes. (a) Volcano plot representing results from the differential expression analysis. Green and purple color indicates genes that are up- and downregulated (FDR ≤ 0.05), respectively, in peripheral blood leukocytes of *Mycobacterium bovis*-infected cattle. Diamond shapes indicate the 272 putatively novel genes found in non-reference sequences. (b) Multidimensional scaling plot of 36 differentially expressed non-reference genes in *Mycobacterium bovis*-infected (blue) and non-infected (orange) Holstein cattle.

Within the 36 putatively novel genes, 28 and 8 are respectively up- and downregulated in peripheral blood leukocytes of *Mycobacterium bovis*-infected cattle, with an average 2-fold change compared to non-infected controls (Supplementary Fig. 9). Multidimensional scaling representations of transcript abundance estimates of the 36 differentially expressed genes separated *Mycobacterium bovis*-infected from non-infected cattle (Fig. 4b). BLASTX queries against a protein reference database provided additional support for 13 out of 36 differentially expressed genes (Supplementary Table 3). The top upregulated non-reference gene supported by the BLASTX query (4.04-fold increase, *P*=1.98e-05) encodes the Workshop Cluster (WC) 1.1-like protein, i.e., a receptor expressed on gamma delta T cells that modulates the immune response to *Mycobacterium bovis* infections^27–29^.

The top downregulated non-reference gene supported by the BLASTX query encodes a protein with high similarity (79.80%) to leukocyte immunoglobulin-like receptor A5 (LILRA5). LILRA5 triggers the strength of the innate immune response to *Mycobacterium* infections^30^ and might serve as a target for pathogen-mediated immunomodulation. Many genes of the leukocyte receptor complex are missing in the assembled chromosomes of the ARS-UCD1.2 reference; instead, *LILRA5* (LOC100139766) is annotated on a 236 kb long unplaced scaffold (NW_020190675). A non-reference gene encoding a protein similar to LILRA5 is located within a 20.4 kb insertion of the multi-assembly graph at 62,471,732 bp on chromosome 18. Both taurine (Original Braunvieh) and indicine (Brahman) assemblies support this insertion. The putatively novel gene encoding LILRA5 is expressed at 9.59±2.54 and 23.10±8.30 CPM, respectively, in *Mycobacterium bovis*-infected and non-infected cattle, corresponding to a 2.19-fold decrease (*P*=1e-04) in infected cattle (Supplementary Table 3).

### Variant discovery from the non-reference sequences

Next, we mapped short sequencing reads, with an average of 19-fold sequencing coverage, from 45 cattle representing five taurine breeds against ARS-UCD1.2 and the extended bovine reference genome. Across 45 samples, the average mapping rate increased by 0.0176% over ARS-UCD1.2, corresponding to approximately 100K sequencing reads for a DNA sample sequenced at 30-fold coverage. The mapping rate increased more noticeably for Brown Swiss (0.024%) and Original Braunvieh (0.021%) than Holstein (0.015%) and Simmental (0.016%) cattle (Supplementary Fig. 10). Similarly, to the transcriptome mapping, sequence reads from Dominette benefitted the least from the extended reference genome (0.006%). However, the increase in mapping rate was greater (0.013%) for other Hereford cattle. For all breeds, the extended reference genome also enabled more perfect alignments (alignments without difference from the reference), less partially mapped (i.e., clipped) reads, and less reads with supplementary alignments. However, the proportion of reads with unique alignment was lower for the extended than standard reference genome (Supplementary Table 4).

We next investigated the alignments against the 2,115,702 non-repetitive non-reference bases detected in all assemblies except ARS-UCD1.2. Among these, 919,761 bases were covered by confident alignments (≥10-fold) from Dominette. This suggests that, although absent from the autosomal assembly, these sequences do occur in the animal used to construct the reference. However, 1,195,941 bp were not covered with reads from Dominette, but instead from Brown Swiss, Holstein, Original Braunvieh or Simmental samples. Strikingly, reads from non-Dominette Hereford samples covered 745,392 of the 1,195,941 bases. This directly implies that Dominette has individual-specific deletions, which are either rare or absent in other Hereford cattle.

Mapping against the extended reference resulted in many reads changing alignment location to the non-reference additions. Most (85.55%) of the reads mapping at non-reference sequences already mapped to the original ARS-UCD1.2 reference genome, although 5% of these mapped to unplaced contigs, while 14.55% were previously unmapped. These mappings displayed an increase in the average mapping quality (22 to 44), alignment score (110 to 142), and alignment identity (0.975 to 0.995). The proportion of clipped reads decreased from 39% to 4%. The subset of these reads which were previously unmapped showed even greater improvements (Supplementary Fig. 11).

Using reads with mapping quality greater than 10 for reference-guided sequence variant genotyping yielded 83,250 filtered variants (73,709 SNPs, 9,541 Indels) in non-reference sequences that were identified by both SAMtools and GATK. These variants formed 80,995 biallelic and 2,255 multi-allelic sites, with a Ti:Tv ratio of 1.91, averaging 1.18 variants per kb. On average each Brown Swiss, Original Braunvieh, Holstein, Simmental, and Hereford animal respectively had 31,028, 29,685, 29,851, 30,309, and 15,845 variant sites in non-reference bases (Fig. 5a). A DNA sample from Dominette had considerably fewer polymorphic sites at non-reference bases, only 7,531. Most variants (32.67%) had alternate allele frequency less than 0.1, and 193 were fixed for the alternate allele (Supplementary Fig. 12). The top principal components from a genomic relationship matrix that was built from the 83,250 non-reference variants separated the animals by breeds (Fig. 5b,c).

**Fig 5:**
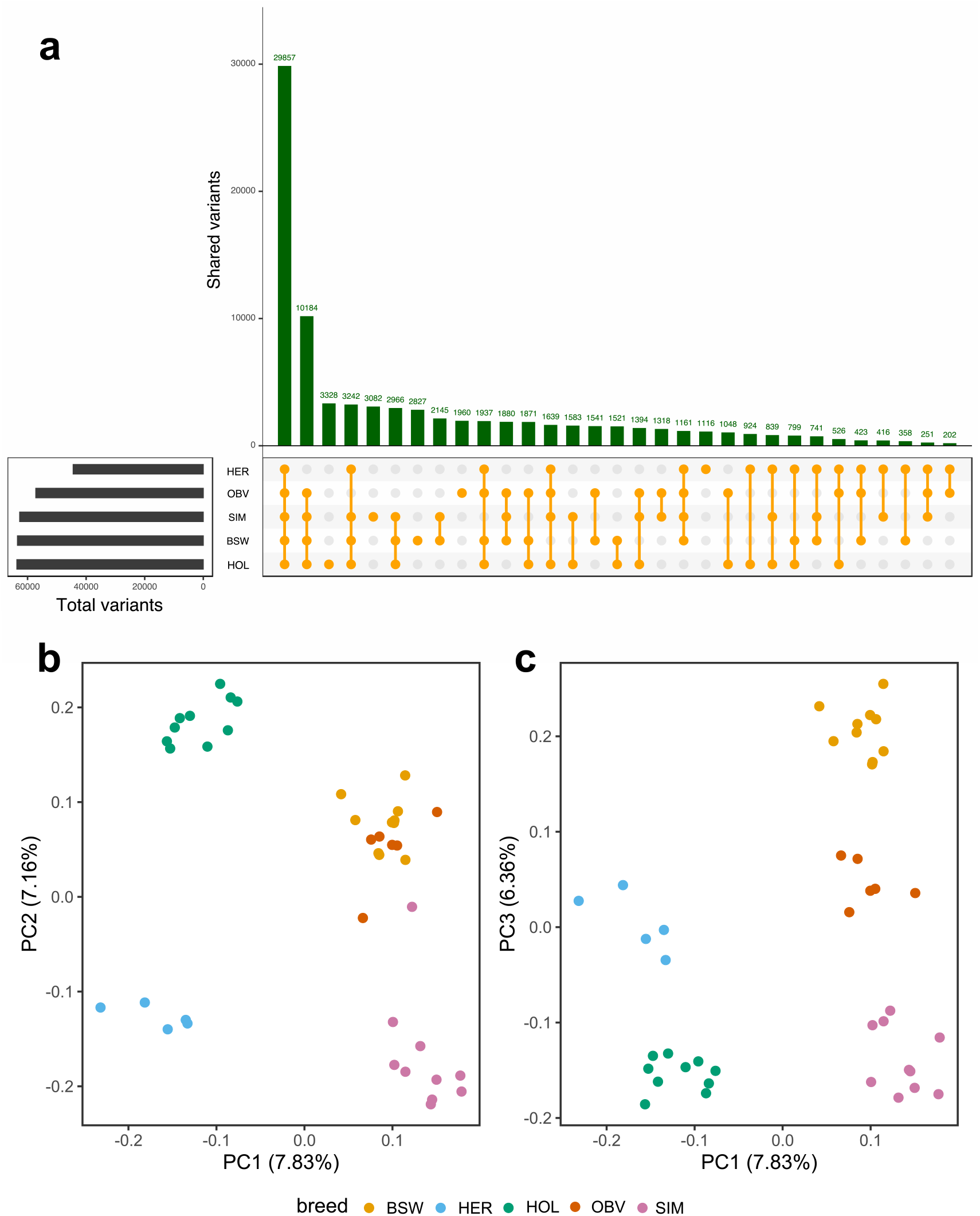
Polymorphic sites detected from non-reference sequences in five breeds. (a) Sharing of 83,250 variants across five taurine cattle breeds (BSW: Brown Swiss, HER: Hereford, HOL: Holstein, OBV: Original Braunvieh, SIM: Simmental). (b, c) The top three principal components (PC) of a genomic relationship matrix constructed from non-reference sequence variants separate the animals by breeds.

## Discussion

We uncover novel non-reference sequences from a bovine multi-assembly graph. Novel contigs can also be assembled from unmapped reads, but placing them onto reference coordinates is difficult if the contigs had been assembled from short reads^12,31^. Our approach provides physical coordinates for the novel sequences because the breakpoints anchor them onto the reference genome. Despite including the genetically distant yak, constructing the multi-assembly graph using minigraph^21^ was computationally efficient and scalable.

Sophisticated algorithms facilitate constructing multi-assembly graphs from thousands of assemblies^32^. To determine the origin of the non-reference sequences, we developed an approach to assign labels to all nodes in the multi-assembly graph. Our evaluation showed that this strategy is highly accurate. Encoding path information in the multi-assembly graph (e.g., with Walk/W-lines) reveals the origin of each node without the need to realign each assembly back to the multi-assembly graph^21^.

We detect 45,357 autosomal segments with a cumulative length of 70,329,827 bases missing in the bovine linear reference genome. To obtain continuous non-reference sequences spanning multiple non-reference nodes, we recovered the non-reference alleles from structural variations. To our knowledge, our study is the first to systematically uncover non-reference sequences in domestic cattle and their close relatives from multiple assemblies. The number of bases missing in the cattle reference genome is comparable to values reported for pigs (72.5 Mb^33^) and goats (38.3 Mb^34^), based on multi-assembly graphs constructed from 11 and 8 animals representing different breeds respectively. In our study, many non-reference sequences originate from yak. Hybridizing between yak and cattle is widely practiced, and results in fertile female descendants. However, multiple generations of backcrossing are required for males to resume fertility^35^. A pangenome constructed from domestic cattle and their extant relatives as recently proposed by the Bovine Pangenome Consortium^36^ will reveal variants that were lost during domestication and the separation of cattle into specialized breeds^37^. For instance, some of the 8 million non-reference bases specific to Brahman might contribute to the adaptation of indicine cattle to harsh environments. Individual taurine assemblies also contain between 14 and 18 million bases that are missing in the Hereford-based reference assembly, many of which are shared between individuals. This value is somewhat higher than the 5-10 million non-reference bases detected per human genome^38–40^, possibly because cattle breeds diverged more strongly than human populations due to intense artificial selection. Each of the three taurine assemblies contains approximately 3 million autosomal non-reference bases that were not detected in any other assembly. There were also 4.4 million non-reference bases, of which 2.1 million were non-repetitive, that were present in all assemblies except the reference. This includes 1.2 million bases that are either specifically deleted in the Hereford breed or the animal used to build the reference, inadvertently propagating reference-bias.

A reference graph may integrate linear reference coordinates, non-reference sequences, and shorter variants^20^. However, as many genome analysis tools still rely on a linear coordinate system, we append the novel non-reference sequences linearly to the ARS-UCD1.2 reference genome. We considered only variations larger than 100 bp because integrating smaller variations increases the complexity of the resulting reference with limited benefit for downstream analyses^21^. We show that our extended bovine reference genome leads to improved DNA and RNA sequence read mapping, even for breeds that did not contribute to the multi-assembly graph. However, excessively adding novel sequences to the reference genome carries the risk of increasing the number of ambiguous alignments.

The non-reference sequences comprise more repetitive elements than the overall ARS-UCD1.2 reference genome (76% versus 48%), but is a lower proportion compared to non-reference insertions detected from human pangenomes (88%)^12,38^. Many non-reference sequences with repetitive elements were observed at bovine immune gene complex loci, corroborating that these regions are highly repetitive^41^. The immune gene complex loci also contain many non-repetitive non-reference sequences suggesting great allelic diversity which may cause assembly problems, thus resulting in gaps and missing sequences in the primary ARS-UCD1.2 assembly.

We show that the 16.6 million non-repetitive non-reference bases encompass transcribed features. An *ab initio* approach predicted 857 gene models from these sequences. The *de novo* assembly of RNA sequencing read alignments from liver samples provided additional support for more than 400 of these gene models. As these analyses were only conducted on liver transcriptomes, it is highly likely that the non-reference sequences contain additional coding sequences that are transcribed in different tissues. The discovery of distinct putatively novel genes in an independent RNA sequencing dataset from peripheral blood leukocytes of Holstein cattle supports this hypothesis. Some of the putatively novel genes, including genes encoding olfactory receptors, were also present in the animal used to build the reference genome. Olfactory receptors have been observed to undergo frequent duplication and rapid evolution in mammalian genomes^42,43^. Segments encompassing duplicated genes may either be collapsed in primary assemblies or result in unplaced contigs that represent variants of the sequence in the assembled chromosomes^44,45^, hence the presence of paralogous copies among non-reference genes is expected. In order to obtain a confident set of non-reference genes, we retained only genes that were not expressed in Dominette. Many of the proteins encoded by these non-reference genes are predicted to play roles in the immune response. Pangenome analyses in species other than cattle have also revealed non-reference genes with immune-related functions^42,46,47^. Our findings show that more novel transcripts can be assembled in breeds that contribute to the multi-assembly graph (Brahman, Angus) than those not included (Holstein, Jersey), suggesting that individual assemblies contain breed-specific, functionally relevant bases. We detect the largest number of non-reference genes using RNA samples from Brahman, suggesting that breeds with great genetic distance from the reference benefit the most from a more diverse reference genome. Importantly, some putatively novel genes are differentially expressed between *Mycobacterium bovis*-infected and non-infected cattle, including genes that encode proteins that either contribute to the immune response against *Mycobacterium* infections or may serve as targets for immunomodulation by the pathogen. These differentially expressed genes remained undetected when the transcriptomes were aligned against the standard bovine reference genome^26^. Thus, our multi-assembly graph uncovers functionally active and biologically relevant genomic features that are missing in the bovine reference genome.

The extended bovine reference genome also leads to substantial improvements over ARS-UCD1.2 in reference-guided alignment and variant discovery. First, the sequence read mapping rate increases for samples from all breeds investigated. Using our extended reference genome would enable mapping approximately 100K previously unmapped reads for samples sequenced at 30-fold coverage. Second, the mapping quality increases for reads that were previously aligned to other positions in ARS-UCD1.2, suggesting that the novel sequences resolve misalignments. These findings agree well with results from species other than cattle, including goats, pigs, and humans^33,34,39^. In addition, we show that the novel sequences contain polymorphic sites that remained hitherto undetected; we discover 83,250 variants that segregate within and between breeds of cattle. A cluster analysis based on these variants separated individuals by breed, suggesting that variable non-reference bases might be associated with breed-specific traits. Thus, our bovine multi-assembly graph makes a previously neglected source of inherited variation amenable to genetic investigations.

The size of the bovine multi-assembly graph will grow as additional reference-quality assemblies from the Bovinae subfamily become available. Assemblies which are more distant will contribute correspondingly to the overall pangenome growth, increasing the flexible part of graph, and reducing the size of the core genome (Supplementary Note 2). In its current implementation, our multi-assembly graph only contains insertions and deletions, as other types of structural variations (e.g., translocations, inversions) that distort the collinearity of the assembly graph cannot be integrated accurately with minigraph. We provide a versatile workflow that facilitates constructing and characterizing multi-assembly graphs for a flexible number of assemblies (https://github.com/AnimalGenomicsETH/bovine-graphs). Our workflow provides tools to determine the origin of non-reference bases, derive structural variations from multi-assembly graphs, predict putatively novel genes and append the novel sequences linearly to a reference genome. We anticipate that the latter will become obsolete as soon as accurate and fast base-level alignment and split-read graph mapping enables the full-suite of genome analyses from a reference graph^48^.

## Methods

### Construction of the multi-assembly graph

We used minigraph^21^ (version 0.12-r389) with option -xggs to integrate six reference-quality genome assemblies into a multi-assembly graph. The current bovine reference genome (*Bos taurus taurus*, ARS-UCD1.2, GCF_002263795.1) and four assemblies that were generated previously are accessible at NCBI: Angus (*Bos taurus taurus*, UOA_Angus_1, GCA_003369685.2)^4^, Brahman (*Bos taurus indicus*, UOA_Brahman_1, GCF_003369695.1)^4^, Highland (*Bos taurus taurus*, ARS_UNL_Btau-highland_paternal_1.0_alt, GCA_009493655.1)^5^, yak (*Bos grunniens*, ARS_UNL_BGru_maternal_1.0_p, GCA_009493645.1)^5^. Additionally, we constructed an assembly from a female Original Braunvieh calf (*Bos taurus taurus*) using PacBio high-fidelity (HiFi) reads (Supplementary Note 1). The sampling of blood from the Original Braunvieh animal and its parents was approved by the veterinary office of the Canton of Zurich (animal experimentation permit ZH 200/19).

The genetic distance among the six assemblies was estimated using Mash (version 2.2)^22^. We performed genomic sketching separately for each assembly with *mash sketch* using a sketch and k-mer size of s=1000 and k = 21, respectively. Sketches were combined using *mash paste*, and *mash dist* was used to estimate the distances between the assemblies. A phylogenetic tree was built from the estimated pairwise distances using the neighbor-joining method^49^ as implemented in the R package ape (version 5.4)^50^. The tree was visualized with the *phylo.plot* function, using the yak assembly as the outgroup to root the tree.

### Identification of non-reference segments from the multi-assembly graph

We refer to nodes that are not in the Hereford-based reference genome (ARS-UCD1.2) as non-reference nodes. We separately aligned (with minigraph parameters “--cov-x asm”) each of the six assemblies back to the multi-assembly graph to determine the support for non-reference nodes. For each alignment, all nodes with non-zero coverage, i.e., nodes traversed by this specific assembly, were labelled. After iterating through all the alignments, each node then contained labels for every assembly which passed through it. As such, each node necessarily had at least one label, while a node traversed by all six assemblies would have six labels (Supplementary Fig. 1).

It was possible to assess minigraph’s alignment accuracy for the path of the Hereford-based reference genome (ARS-UCD1.2), because all reference nodes in the multi-assembly graph were from this assembly. Nodes were considered true positive (TP) and true negative (TN) when reference and non-reference nodes were correctly assigned Hereford labels, respectively. Reference nodes aligned as non-reference nodes were assigned false negative (FN) and non-reference nodes aligned as reference nodes were assigned false positive (FP). We characterized alignment recall (TP / (TP+FN)), precision (TP / (TP+FP)), and overall F1 score (2 * (precision * recall) / (precision + recall)).

### Identification of structural variations from the multi-assembly graph

We used the bubble popping algorithm of gfatools (version 0.4)^21^ to derive the structural variations from the multi-assembly graph. In the reference graph model of minigraph, a bubble is a branching region in the graph for which the start and end node are reference sequences. A path traversing the start and end nodes represents an allele of a structural variant.

The version of gfatools considered in our study reports the shortest and longest path for each bubble. To detect and classify all paths within a bubble, we applied the following stepwise procedure (Supplementary Fig 2):

- Determine the start and stop node for each bubble using the bubble popping algorithm of gfatools.
- Traverse all possible paths in the bubble using a recursive depth-first search.
- Retain only paths with color-consistent labels (see above).
- Classify a path as a reference path when all nodes and edges are part of the Hereford-based reference assembly, and as non-reference otherwise.
- Compare reference and non-reference paths to classify the type of the structural variations.

Structural variations were classified as biallelic if two paths were observed in a bubble and multi-allelic if a bubble contained more than two paths. The structural variations were further classified into:

- Alternate deletion, when the non-reference path was shorter than the reference path (but the reference path has nonzero length).
- Complete deletion, when the non-reference path has a length of zero.
- Alternate insertion, when the non-reference path was longer than the reference path.
- Complete insertion, when the reference path has a length of zero.

Breakpoints of structural variations were determined according to ARS-UCD1.2 reference coordinates. We overlapped the breakpoints with annotations from Ensembl (build 101) to identify structural variations in coding sequences. Affected genes were subjected to a gene set enrichment analysis using PANTHER (http://pantherdb.org/)^23^ for which the *Bos taurus* reference gene list was supplied as a baseline.

To validate the structural variations, we mapped 6,803,270 (~46-fold coverage) PacBio HiFi reads to the multi-assembly graph using GraphAligner (version 1.0.12)^24^ with preset-x vg (variation graph mapping). The HiFi reads were generated from a Nelore x Brown Swiss crossbred bull (SAMEA7765441), representing taurine and indicine breeds that were not used to build the multi-assembly graph. The veterinary office of the Canton of Zurich approved the sampling of blood from the crossbred animal and its parents (animal experimentation permit ZH 200/19). The mean read length was 20,612 bases with an average accuracy of 99.76%. We calculated coverage (number of reads aligned) at each node and edge in the graph based on the GAF (Graphical Alignment Format) output from GraphAligner.

We combined all non-reference alleles (excluding complete deletions, paths without non-reference bases, and paths with length less than 100 bp) to obtain a comprehensive set of non-reference bases from the multi-assembly graph. To facilitate the mapping of short reads to the segment edges, we added 100 bp of flanking sequences (derived from sequences at the source and sink nodes) on either side of the structural variations. The flanking sequences were not considered for length calculations or gene predictions (see below).

To investigate the repeat content of the non-reference sequences, we used the RMBlastn search engine (version 2.10.0) to run RepeatMasker version 4.1.1 (option-species cow)^51^ using the database of repetitive DNA elements from Repbase (release 20181026)^52^.

### Bioinformatic characterization of non-reference sequences

In order to reveal functionally active non-reference sequences, we performed two complementary analyses:

First, we compared the repeat masked non-reference sequences against a local protein database using DIAMOND BLASTX (version 0.9.30)^53^. Using DIAMOND makedb, the local protein database was built from the RefSeq protein sequences of

- Taurine cattle (*Bos taurus taurus*, GCF_002263795.1_ARS-UCD1.2_protein.faa)
- Indicine cattle (*Bos taurus indicus*, GCF_003369695.1_UOA_Brahman_1_protein.faa)
- Yak (*Bos mutus*, GCF_000298355.1_BosGru_v2.0_protein.faa)
- Human (*Homo sapiens*, GCF_000001405.39_GRCh38.p13_protein.faa)
- Mouse (*Mus musculus*, GCF_000001635.26_GRCm38.p6_protein.faa)
- Bison (*Bison bison*, GCF_000754665.1_Bison_UMD1.0_protein.faa)
- Water buffalo (*Bubalus bubalis*, GCF_003121395.1_ASM312139v1_protein.faa)
- Goat (*Capra hircus*, GCF_001704415.1_ARS1_protein.faa)
- Sheep (*Ovis aries*, GCF_002742125.1_Oar_rambouillet_v1.0_protein.faa)
- the curated protein databases of SwissProt and PDB (ftp://ftp.ncbi.nlm.nih.gov/blast/db/FASTA/)

To query the non-reference sequences against the local protein database we ran BLASTX with the parameters “--more-sensitive --e-value 1e-10 --outfmt 6”. We considered only the top hit for each queried sequence with minimum coverage and identity of 80%.

Second, we performed an *ab initio* gene structure prediction from the repeat masked non-reference sequences using a local instance of Augustus (version 3.3.3)^54^ using parameters trained on the human genome. From the Augustus GTF output file, we extracted the number of gene models, the number of gene models with transcription start and termination site, transcript length, exon count, and length per gene, coding sequence count and length per gene, and protein length of the putatively novel protein-coding sequences. To classify the domain and family of the putatively novel proteins, we converted the Augustus GTF output to the fasta format and performed a query against the local protein database (as above) using DIAMOND BLASTP with the same parameters and thresholds as the BLASTX query.

### *De novo* transcript assembly from non-reference sequences

We downloaded between 12,361,440 and 34,421,106 paired-end RNA-sequencing reads from liver tissue from 10 Angus^55^, 10 Brahman^56^, 9 Holstein^57^ and 10 Jersey^57^ cattle, as well as from Dominette -- the animal used to construct the ARS-UCD1.2 reference^2^). Adapter sequences and low-quality bases were removed from the raw RNA sequencing data using default parameters of fastp (version 0.19.4)^58^. The filtered reads were then aligned using HISAT2 (version 2.1.0)^59^, with option “--dta” to facilitate the downstream transcriptome assembly, to the original ARS-UCD1.2 reference as well as the extended version of the ARS-UCD1.2 reference. The extended reference was constructed by appending repeat masked non-reference sequences as unplaced contigs^59^.

Putative novel transcripts were assembled *de novo* using StringTie2 (version 2.1.1) ^25^ from RNA-seq reads that aligned to the non-reference sequences^25^. To facilitate transcript assembly, we supplied the ARS-UCD1.2 Ensembl annotation (build 101) and the novel gene models predicted by Augustus (see above). Transcripts were assembled *de novo* separately for all RNA sequencing samples. Subsequently, we used StringTie2 *merge* to create a unique set of transcripts across all samples and facilitate the assembly of full-length transcripts from partially assembled transcripts. We quantified gene expression for each sample with StringTie2 using a fixed (merged) GTF file that was generated previously (without predicting novel transcripts, option -e). Gene abundance was quantified in transcript per million (TPM).

### Differential gene expression analysis

We utilized publicly available peripheral blood leukocyte transcriptomes of eight *Mycobacterium bovis*-infected and eight age-matched healthy Holstein cattle^26^ to detect differentially expressed genes from non-reference sequences. The RNA-sequencing data contain between 9,272,629 and 25,358,979 single-end reads of length 78 bp. We performed quality control on the raw sequencing reads using fastp (version 0.19.4)^58^ with default parameters. The filtered reads were then mapped to the extended ARS-UCD1.2 reference genome that contained the non-reference sequences using HISAT2^59^. Potential non-reference transcripts were assembled *de novo* with StringTie2 (see above). Gene-level read counts were estimated based on a custom annotation file that contained the Ensembl (build 101) ARS-UCD1.2 genome annotation and the non-reference annotation as generated by StringTie2 using the *featurecounts* function of the Rsubread package (option countMultiMappingReads =FALSE to exclude multi-mapping reads). The read count matrix was used as input for EdgeR version 3.24.3^60^. We normalized transcript abundance by sequencing depth using the trimmed-mean of M-values (TMM) approach. Genes that were expressed at ≥1 count per million (CPM) in at least eight samples were tested for differential expression in peripheral blood leukocytes between *Mycobacterium bovis*-infected and control animals using a generalized linear model (GLMQfit) with dispersion parameter estimated using the Cox-Reid method. Genes were considered to be differentially expressed at a Benjamini-Hochberg-corrected FDR≤0.05. Multidimensional scaling of the normalized read count matrix of the differentially expressed genes was performed using the *cmdscale* function in R.

### Mapping and variant calling from whole-genome short read data

We considered the original ARS-UCD1.2 reference genome and an extended version of the reference that additionally contained 70,329,827 non-reference bases detected from five assemblies. We used paired-end short read sequencing data from 45 samples representing five breeds: OBV, BSW, Holstein, Simmental^61^, and Hereford (including Dominette, the animal used to construct the ARS-UCD1.2 reference genome)^2,62^ that had average sequencing coverage of 18.94-fold. Quality control of the short-read sequencing reads was performed using fastp (version 0.19.4)^58^ with default parameter settings. The filtered reads were subsequently mapped to the original ARS-UCD1.2 reference and the extended ARS-UCD1.2 reference that also contained non-reference sequences using the mem-algorithm of BWA (version 0.7.17)^63^ with default parameters. Duplicate reads were marked with Samblaster (version 0.1.24)^64^.

We performed multi-sample variant calling (SNP and Indels) on the non-reference sequences using SAMtools (version 1.10)^65^ and GATK (version v4.1.9.0)^66^ as detailed in Crysnanto *et al.*^17^. Base quality scores were recalibrated using known variants from the 1000 bull genomes project database (http://www.1000bullgenomes.com/doco/ARS1.2PlusY_BQSR_v3.vcf.gz). We applied the GATK modules *HaplotypeCaller*, *GenomicsDBImport* and *GenotypeGVCFs* to discover and genotype polymorphic sites. The variants were subsequently hard-filtered using recommended parameters (SNP filters: QD < 2 || QUAL < 30 || FS > 60 || MQ < 40 || MQRankSum < −12.5 || ReadPosRankSum < −8 || AN < 10, Indel filters: QD < 2 || QUAL < 30 || FS > 200 || ReadPosRankSum < −20.0 || AN < 10)^17^. A second independent variant discovery and genotyping approach was performed using SAMtools *mpileup* and bcftools *call*^65^. The resulting genotypes were subsequently hard-filtered according to parameters recommend by the 1000 bulls genomes project (QUAL < 20 || MQ < 30 || DP < 10 || AN < 10)^7^. To create a consistent variant representation across both datasets, variants were normalized using vt version 0.5^67^. We retained only filtered variants, which were identified by both SAMtools and GATK.

## Data availability

Short sequencing reads are available at the European Nucleotide Archive (ENA) (http://www.ebi.ac.uk/ena) with study accession PRJNA436715 (Transcriptome - Brahman), PRJNA392196 (Transcriptome - Angus), PRJNA357463 (Transcriptome – Holstein, Jersey), PRJNA294306 (Transcriptome-Dominette), PRJNA257841 (Differential expression analysis – Holstein), PRJEB18113 (WGS – BSW, OBV, HOL, SIM), PRJNA494431 (WGS - Hereford), PRJNA391427 (WGS - Dominette). Accession numbers for all samples are provided in Supplementary Data 1. PacBio HiFi reads for an Original Braunvieh animal used to construct a *de novo* assembly are available at study accession PRJEB42335 under sample accession SAMEA7759028. PacBio HiFi reads for a Nelore x Brown Swiss bull are available at study accession PRJEB42335 under sample accession SAMEA7765441. Data supporting this study, including the multi-assembly graph, non-reference sequences, putatively novel genes, transcript abundances and sequence variants detected from non-reference sequences are available via Zenodo (https://doi.org/10.5281/zenodo.4385983)^68^.

## Code availability

Workflows to construct multi-assembly graphs and custom scripts to characterize non-reference sequences are available via Github (https://github.com/AnimalGenomicsETH/bovine-graphs). All workflows were built using Snakemake (version 5.30.1)^69^ and custom scripts were written in R (version 3.5.1)^70^ and Python (version 3.7.1).

## Acknowledgements

We are thankful for the excellent technical support provided by the ETH Zürich functional genomics platform FGCZ (https://fgcz.ch/). Computing was done at the Leonhard High Performance Compute cluster at ETH Zürich. This study was supported by grants from the Swiss National Science Foundation (310030_185229) and the Swiss Federal Office for Agriculture (FOAG), Bern.

## Author contributions

HP and DC conceived the study; DC and HP designed the analyses and DC and AL performed them; ZHF coordinated the sampling and long-read sequencing; AL assembled the Original Braunvieh genome; DC and HP prepared the manuscript with all authors contributing to the final version of the paper.

