## Supplementary Information for "Novel functional sequences uncovered through a bovine multi-assembly graph"

Danang Crysnanto*, Alexander S. Leonard, Zih-Hua Fang, Hubert Pausch
Animal Genomics, ETH Zürich, Zürich, Switzerland

#### Supplementary Information

Supplementary Figure S1-S12
Supplementary Table S1-S4
Supplementary Note S1-S3

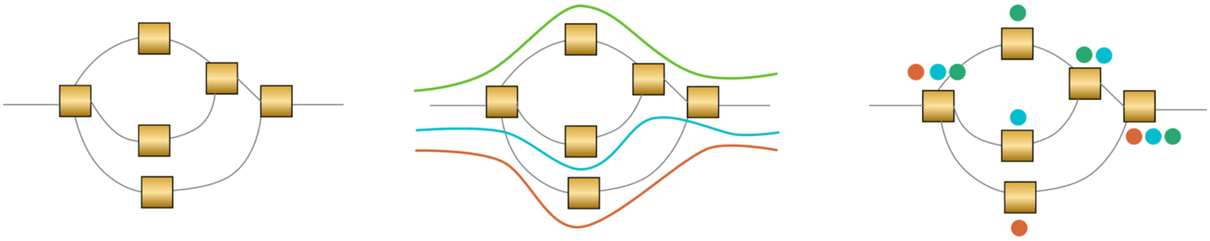

**Supplementary Figure 1. Labelling of the nodes in the multi-assembly graph**. To determine the support for the nodes in the graph, we aligned each individual assembly back to the multi-assembly graph and labeled nodes according to the assembly paths that traversed them with different colors. The left panel represents a schematic graph. Rectangles and lines represent nodes and edges, respectively. The middle panel represents the paths of three assemblies traversing the nodes. The right panel displays how each node that was traversed by an assembly receives a label (colored dots).

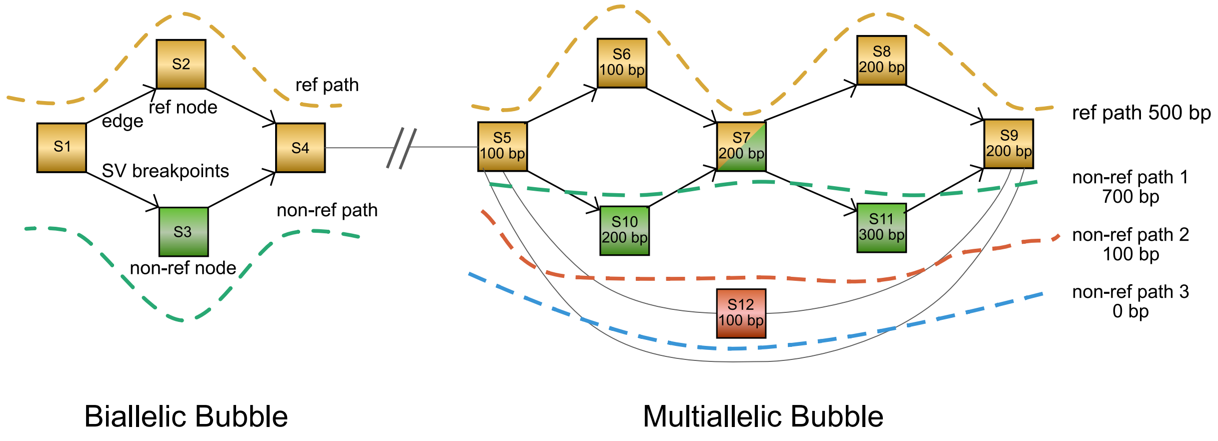


**Supplementary Figure 2. Graphs and structural variants terminology used in the paper**. (*left*) A node contains a sequence of nucleotides (S1-S12). Reference nodes (S1, S2, S4) are derived from the backbone assembly used to construct the graph. Non-reference nodes (S3) contain sequences from additional assemblies that are not present in the backbone. Nodes are connected by directed edges from parent to child where the underlying sequences are contiguous. Edges between reference and non-reference nodes are breakpoints of structural variations. Bubbles are branching regions in the graph which start and end at reference nodes. (*right*) Paths in the bubbles represent different alleles of structural variations, which are biallelic if a bubble contains two paths or multiallelic if it contains more. Nodes within biallelic bubbles represent alleles. Within multi-allelic bubbles, multiple nodes may be part of the same path and thus allele. It is worth noting that not all combinations of nodes within bubbles are real paths found in individual assemblies (e.g., S10-S7-S8). As such, color-consistent nodes within a bubble are stitched together to represent true paths. By comparing reference and non-reference paths, it is possible to determine the type of the structural variations (e.g., non-ref path 1: alternate insertion, path 2: alternate deletion, path 3: complete deletion).

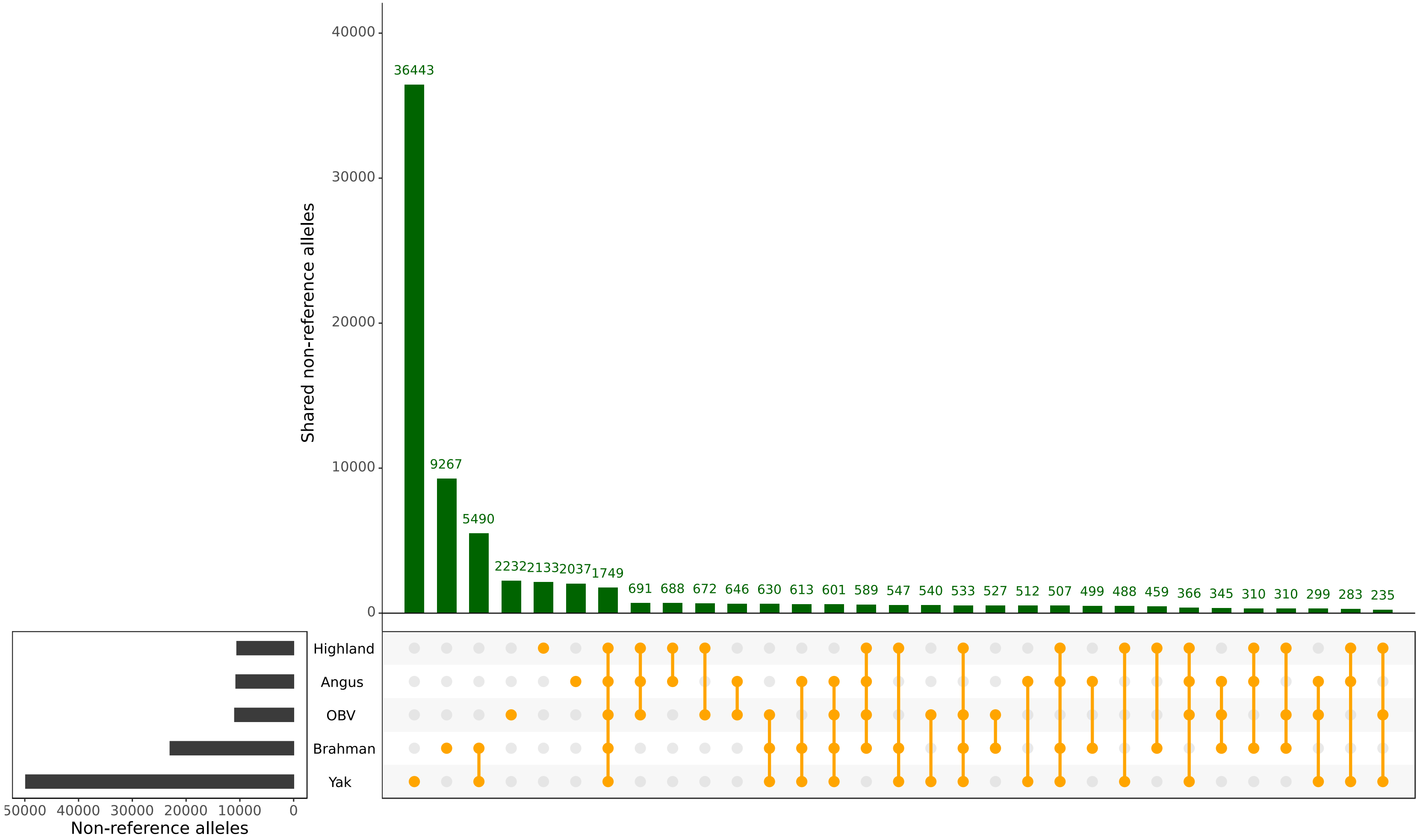


**Supplementary Figure 3. The sharing of 73,506 non-reference alleles across five assemblies in the multi-assembly graph.**

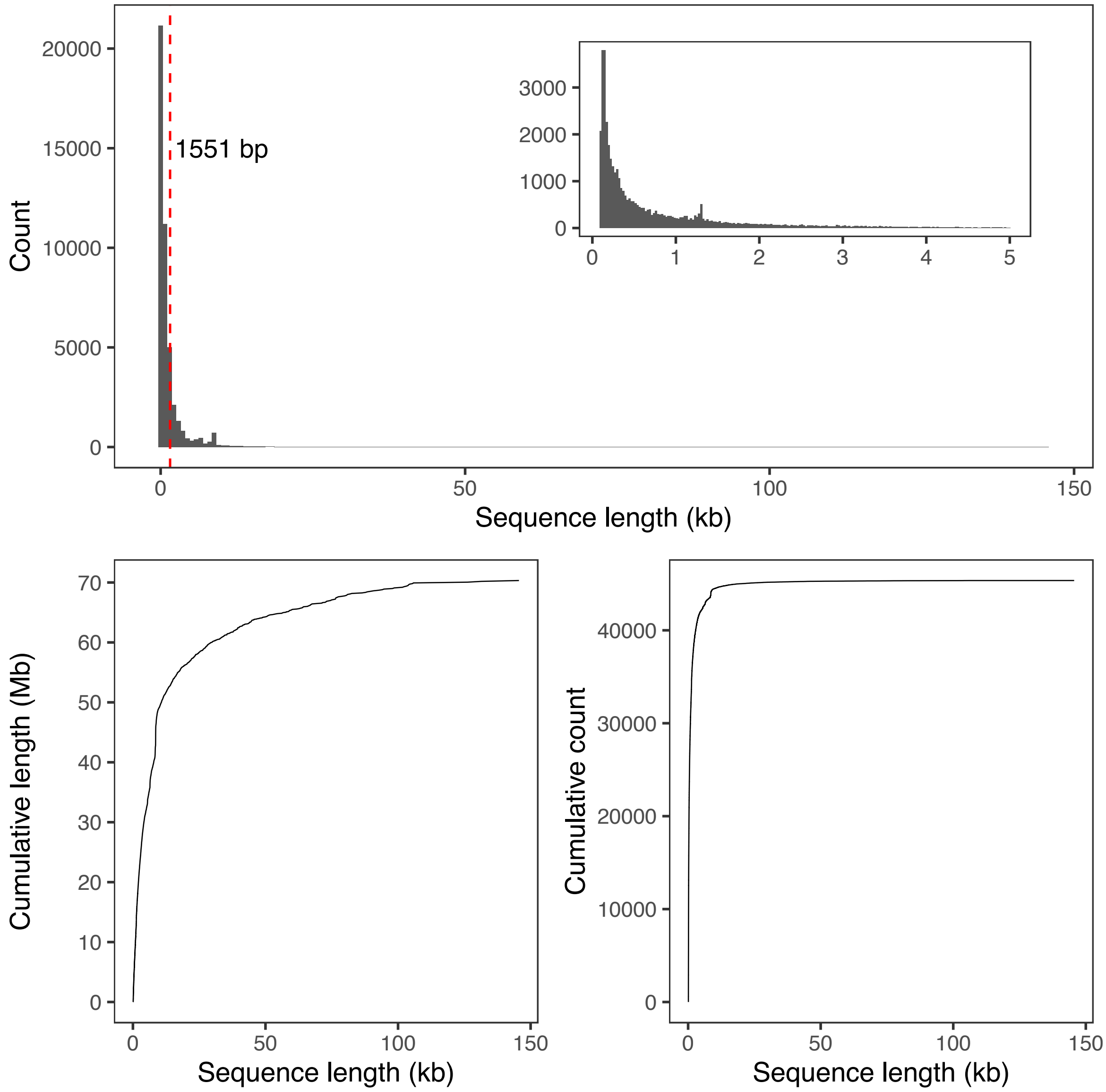


**Supplementary Figure 4. Length of the non-reference sequences that were added linearly to the ARS-UCD1.2 reference**. Length distribution of the non-reference alleles (upper panel) and their cumulative length and count (lower panels). The inset in the upper panel displays the distribution of non-reference alleles shorter than 5 kb. The dashed-red line indicates the average length (1551 bp) of the non-reference alleles.

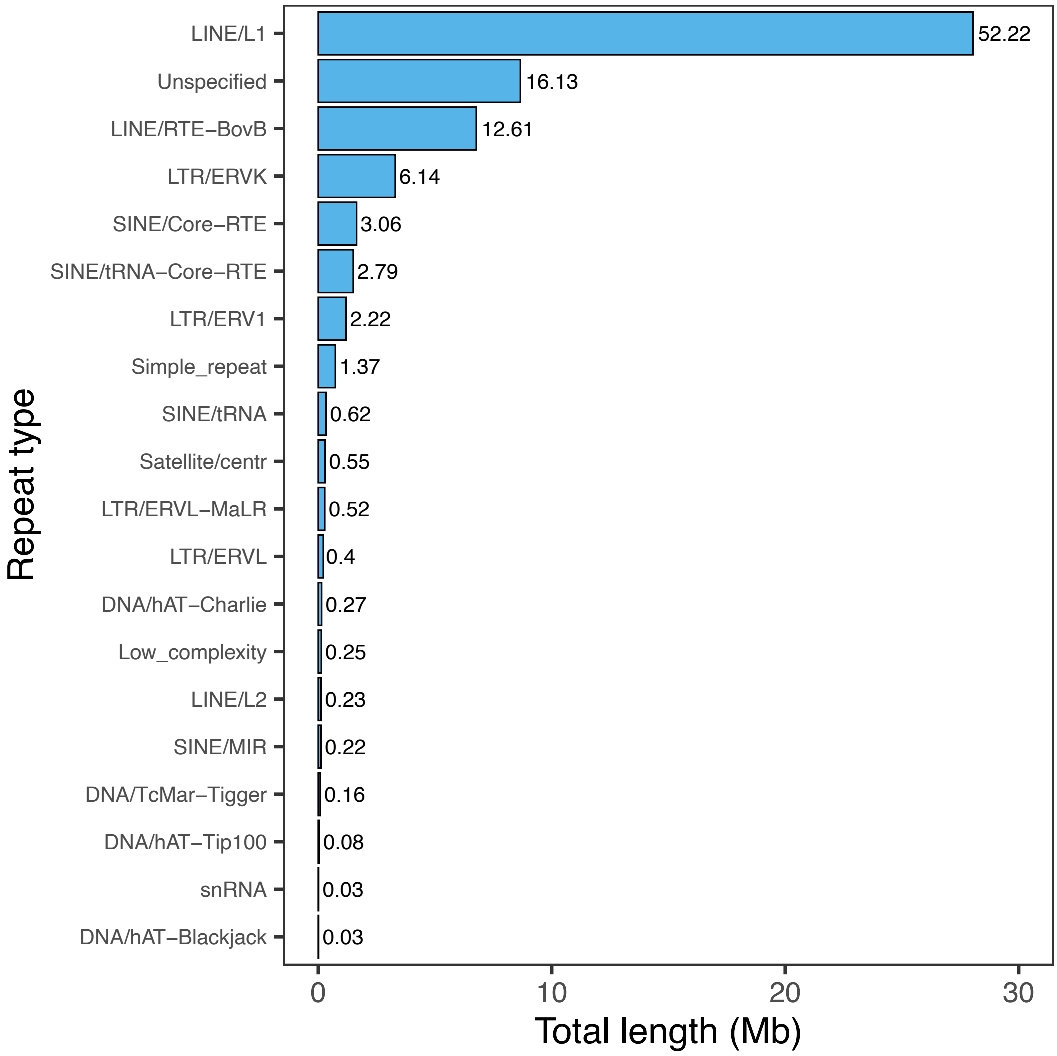


**Supplementary Figure 5. Prevalence of repetitive elements in the non-reference sequences.** The 20 most prevalent repetitive elements account for 99.9% of the repetitive elements detected in the non-reference sequences. The X-axis indicates the summed sequence length (in Mb) spanned by the repetitive elements, with text labels indicate the proportion (%) of a repetitive element contributing to the total repeat length.

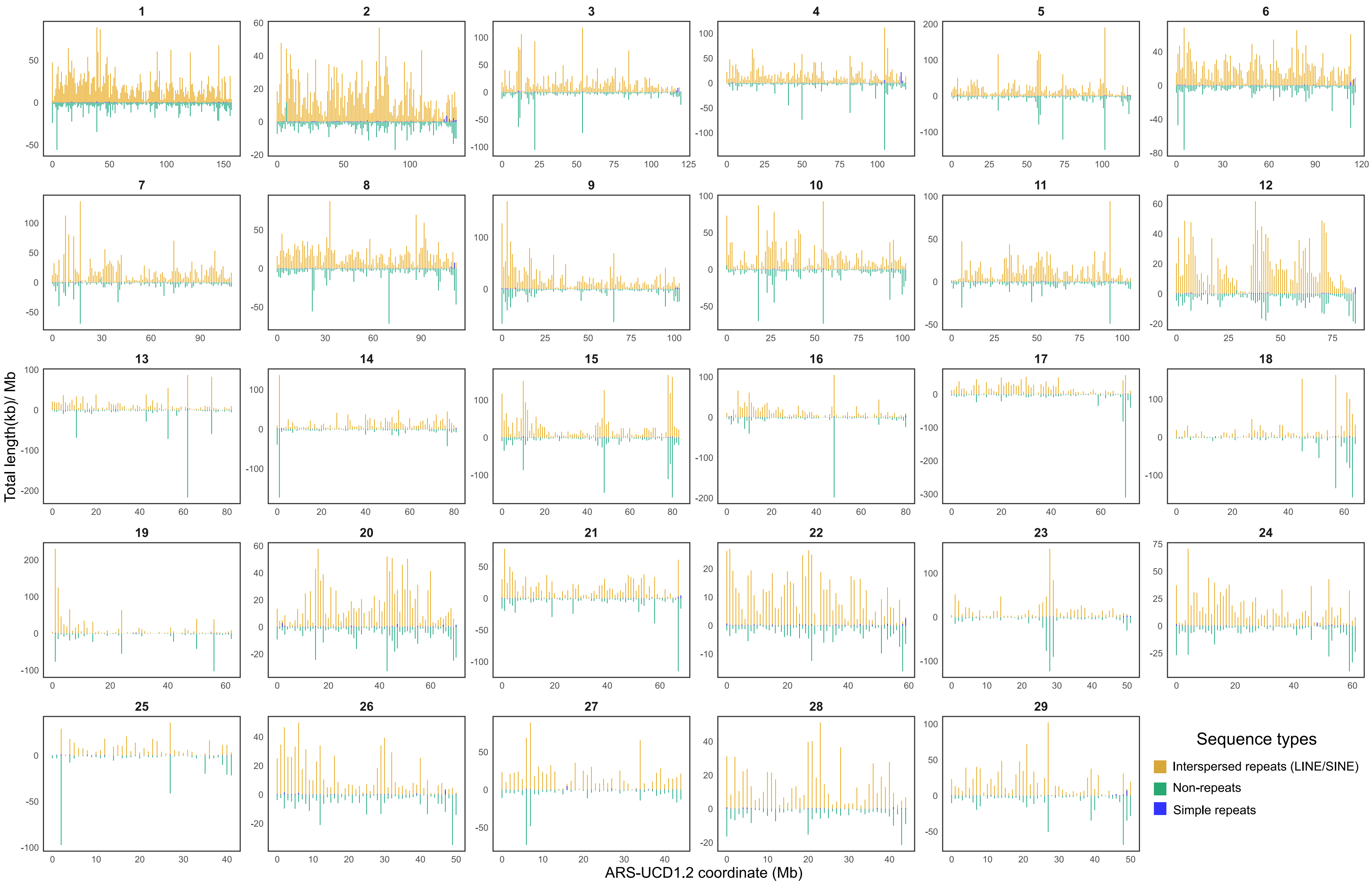


**Supplementary Figure 6.** The distribution of interspersed and simple repeats, and non-repetitive elements found in non-reference sequences based on the ARS-UCD1.2 coordinate system. To aid visualization, the distribution of non-repetitive segments is mirrored to the negative Y-axis. The numbers above the individual panels are chromosome identifiers.

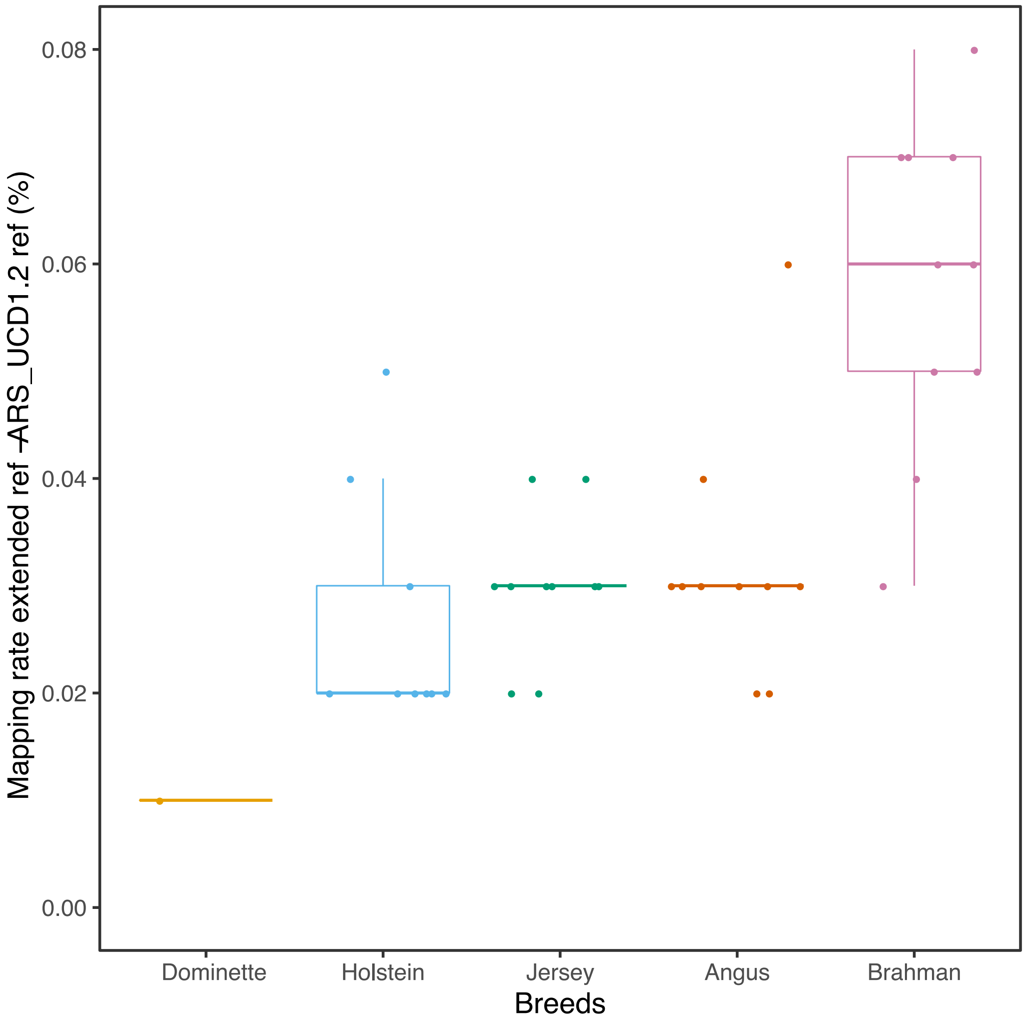


**Supplementary Figure 7. Transcriptome mapping rate improvements in five breeds using the extended reference sequence over ARS-UCD1.2.** Values along the Y axis represent the difference in mapping rate between the extended and the original ARS-UCD1.2 reference (%) as reported by HISAT2. Positive values indicate that more reads aligned to the extended than original reference. Dominette is a Hereford animal used to construct ARS-UCD1.2.

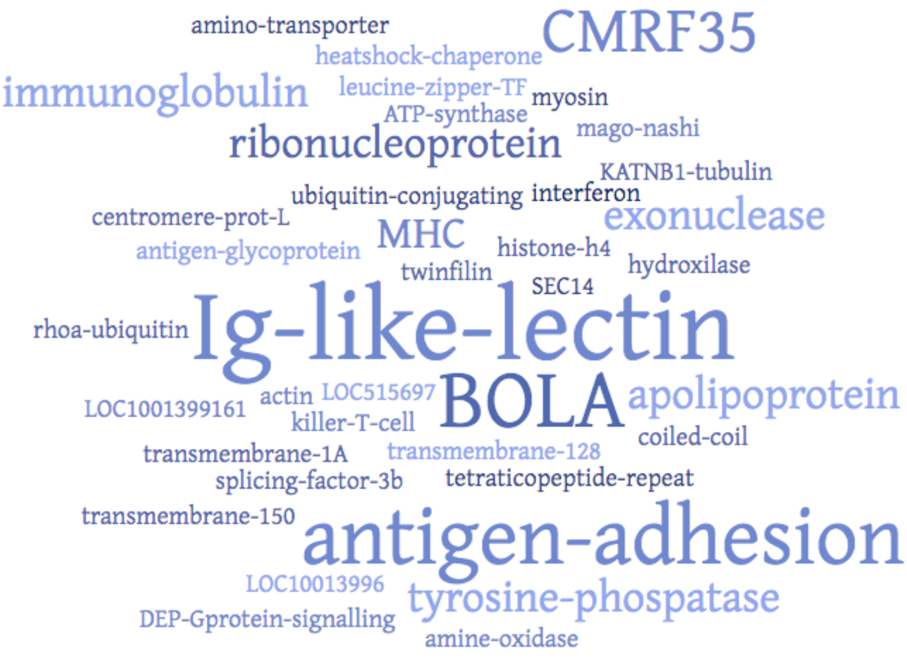


**Supplementary Figure 8. Word cloud of the top blast hits from 142 putatively novel genes assembled from RNA sequencing reads mapping to non-reference sequences.** The BLAST query was performed against a protein database containing sequences from *Bos* and related species. Word size reflects the frequency of the hits.

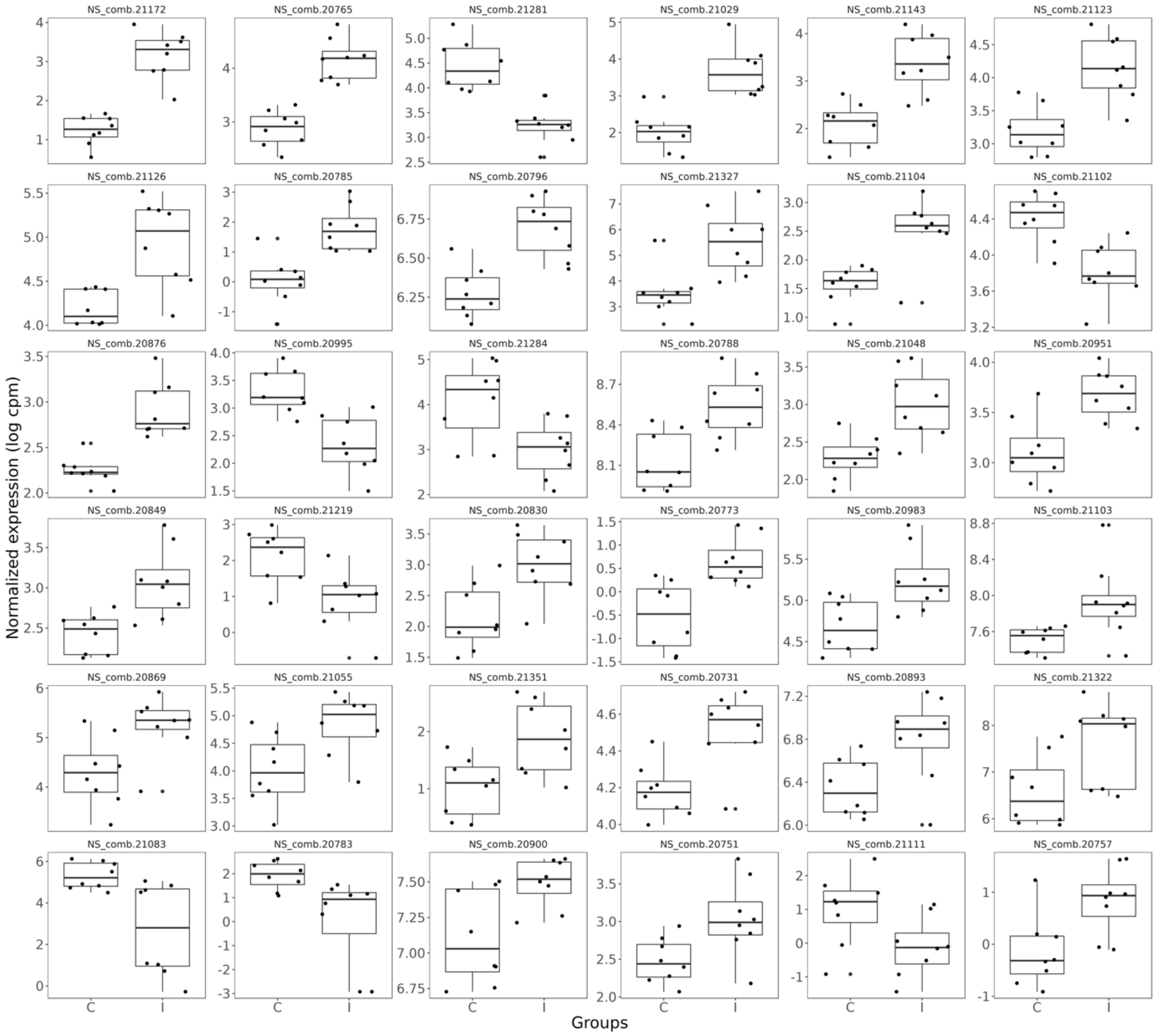


**Supplementary Figure 9. Differential expression of 36 non-reference genes in *Mycobacterium bovis*-infected cattle**. Control (C) and *Mycobacterium bovis*-infected (I) cattle are grouped separately for each gene. *Y* axis indicates the normalized transcript abundance expressed as log2 CPM as reported by EdgeR.

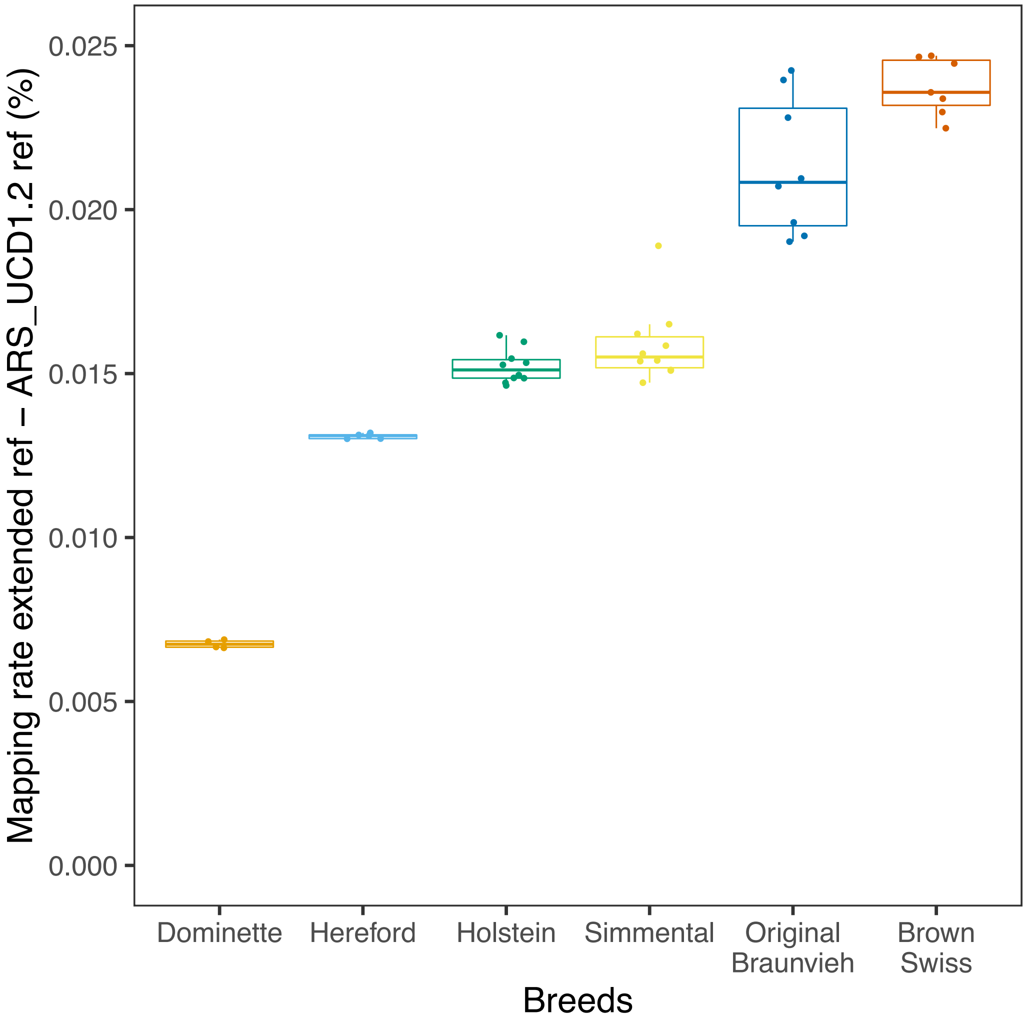


**Supplementary Figure 10. Mapping rate of whole-genome short sequencing reads to the extended linear reference genome.** The Y-axis reflects the difference (in %) in mapping rate between the extended reference and the original ARS–UCD1.2 reference sequences. Positive values indicate that the mapping rate is higher for samples aligned to the extended than original ARS-UCD1.2 reference sequences. Short sequencing reads of 45 cattle from five breeds were considered. Dominette is a Hereford cattle, but is separated as she is the animal used to construct ARS-UCD1.2.

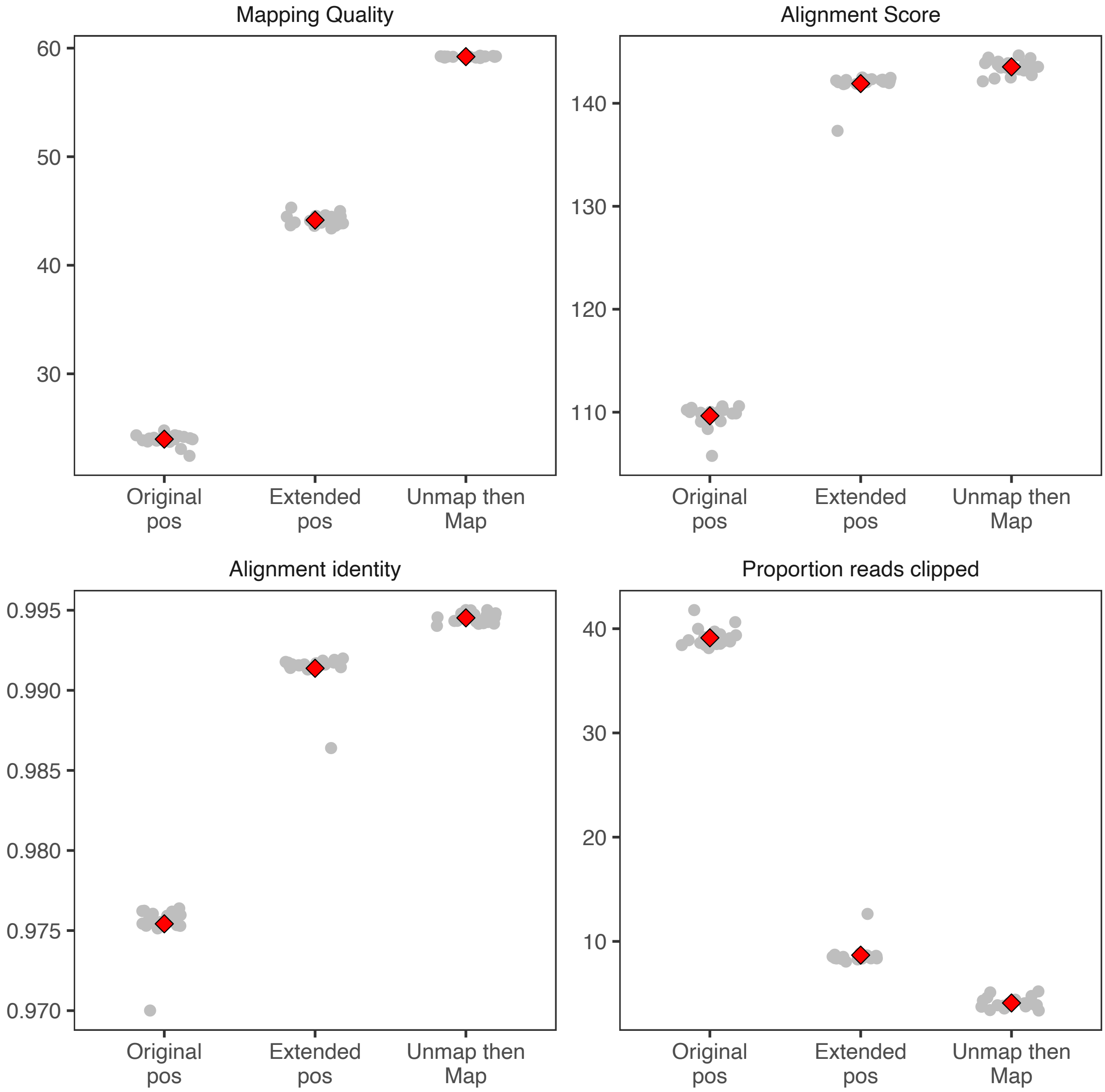


**Supplementary Figure 11. Accuracy of read mapping to non-reference sequences.** Four mapping statistics (mapping quality, alignment score, alignment identity, proportion of clipped reads) were assessed for short sequencing reads from 45 samples across 5 breeds. First, we consider reads that mapped to autosomal sequences of the ARS-UCD1.2. The mapping statistics of these reads are compared between the ARS-UCD1.2 reference sequence (Original pos) and their mapping position at the novel non-reference sequences of the extended reference genome (Extended pos). Second, we consider reads that were unmapped against the ARS-UCD1.2 reference genome but received a mapping position against the extended reference genome (Unmap then map). Each grey point indicates the average mapping statistics for one DNA sample and red diamond indicates the average across all animals.

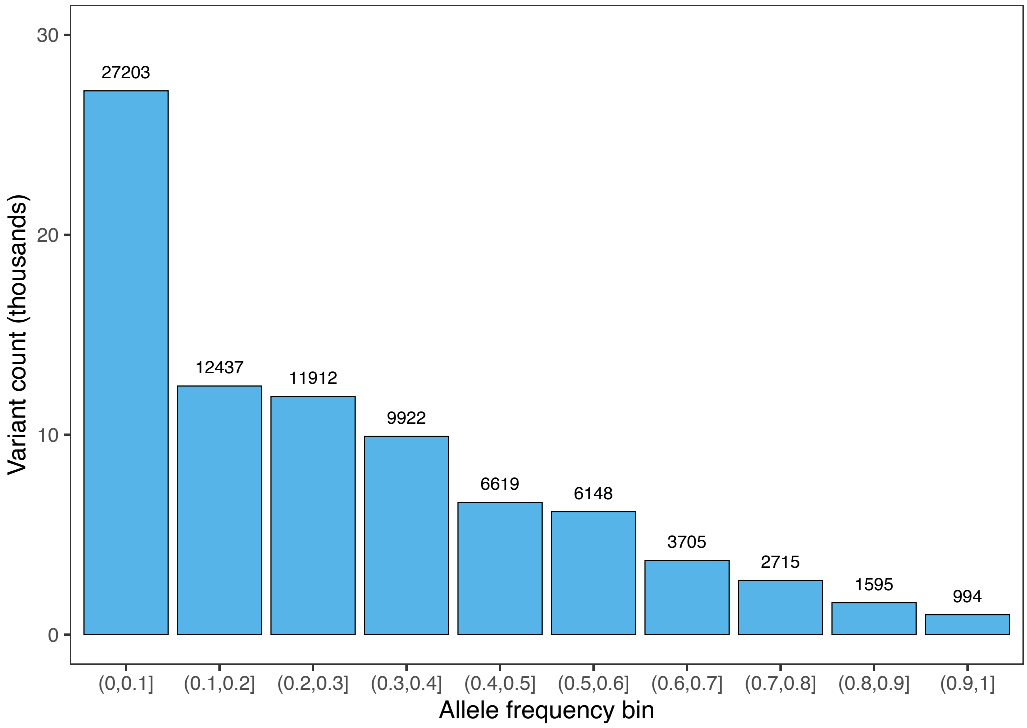


**Supplementary Figure 12. Alternate allele frequency of 83,250 variants detected from non-reference sequences in 45 samples from 5 breeds.**

**Supplementary Table 1. Different types of structural variations discovered from the multi-assembly graph.** Variant length is calculated based on the absolute difference between reference and non-reference allele.

| Mutations | Types | Count | Complete type | Alternate type | Non-ref allele length | Variant length |
|---|---|---|---|---|---|---|
| Insertions | biallelic | 35748 | 20432 | 15316 | 40361474 | 37388222 |
| Insertions | multiallelic | 4621 | 4221 | 400 | 21116534 | 10303720 |
| Deletions | biallelic | 28476 | 15377 | 13099 | 2845080 | 28373582 |
| Deletions | multiallelic | 4661 | 1972 | 2689 | 10130841 | 11727721 |
| ***Total*** | | ***73506*** | ***42002*** | ***31504*** | ***74453929*** | ***87793245*** |

**Supplementary Table 2. Gene model prediction from repeat masked non-reference sequences.** Total novel genes denote all gene models (including partial genes) predicted by Augustus. Complete gene models restricted to only full gene models (TSS, start codon, exon, intron, stop-codon present). Transcript, exon, and CDS statistics reported as mean (maximum-minimum) length from the full gene models.

| Feature of the gene model | Value |
|---|---|
| Total novel genes (distinct SVs) | 857 (768) |
| Complete novel genes (distinct SVs) | 374 (328) |
| Transcript length (bp) | 4742.14 (min: 314; max: 104024) |
| Exon length (bp) | 942.30 (min: 15; max: 6725) |
| Exon length/gene (bp) | 2050.89 (min: 314; max: 7762) |
| Exon count/gene (bp) | 2.18 (min: 1; max: 20) |
| CDS length (bp) | 396.64 (min: 5; max: 3059) |
| CDS length/gene (bp) | 794.34 (min: 199; max: 6280) |
| protein length (aa) | 264.78 (min: 66.33; max: 2093.33) |

**Supplementary Table 3. BLASTX hits of the transcripts from differentially expressed non-reference genes.** Log2 FC is the difference in expression between *Mycobacterium bovis*-infected and non-infected control cattle (e.g., a positive value indicates that expression is higher in infected than control cattle), and Adj FDR is the adjusted false discovery rate determined using the Benjamini-Hochberg correction.

| **Hits** | **Mean (SD) expression in CPM** | | **log2 FC** | **Adj FDR** |
|---|---|---|---|---|
| | Control | Infected | | |
| Antigen WC1.1-like | 2.43 (0.6) | 9.54 (3.65) | 2.0137 | 1.98E-05 |
| Leukocyte immunoglobulin-like receptor subfamily A member 5 isoform X1 | 23.10 (8.30) | 9.59 (2.54) | -1.2870 | 0.0001 |
| PREDICTED: synaptobrevin homolog YKT6 | 4.39 (1.36) | 11.19 (4.7) | 1.3754 | 0.0008 |
| PREDICTED: major vault protein isoform X1 | 21.70 (3.87) | 14.23 (2.88) | -0.6140 | 0.0040 |
| PREDICTED: heat shock 70 kDa protein 1B | 10.18 (2.7) | 5.33 (1.94) | -0.9511 | 0.0041 |
| Elongation factor 1-alpha 1 | 282.86 (40.74) | 374.22 (63.08) | 0.4033 | 0.0093 |
| Heterogeneous nuclear ribonucleoprotein R isoform 2 | 5.47 (0.94) | 8.65 (2.73) | 0.6740 | 0.0148 |
| PREDICTED: prothymosin alpha isoform X2 | 22.02 (10.73 | 39.85 (12.84 | 0.8523 | 0.0243 |
| Stathmin isoform a | 17.56 (7.39) | 30.3 (10.17) | 0.7823 | 0.0271 |
| Serine/arginine repetitive matrix protein 1 isoform X1 | 18.3 (1.74) | 22.99 (3.29) | 0.3293 | 0.0285 |
| BOLA class I histocompatibility antigen, alpha chain BL3-7-like | 109.32 (61.3) | 223.94 (116.59) | 1.0387 | 0.0302 |
| Predicted gene, EG665562 | 141.66 (31.9) | 179.9 (20.92) | 0.3440 | 0.0400 |
| PREDICTED: GTP-binding protein SAR1a | 5.71 (1.22) | 8.71 (3.28) | 0.6231 | 0.0415 |

**Supplementary Table 4. Comparison of read mapping accuracy between the extended and ARS-UCD1.2 reference.** All metrics were extracted from BAM files using pysam v0.16.0.1[1]. Alignment identity reflects the proportion of bases from an aligned read that match the reference sequence. A read was considered to be clipped if the CIGAR string of the alignment contains tags for either hard- (H) or soft-clipped (S) bases. Supplementary alignments were reported for alignments with an XS tag. Criteria for perfect and unique alignments were based on those reported by Crysnanto and Pausch[2]. Specifically, reads were considered to align perfectly if the edit distance was zero along the entire read (NM:0 tag), and when the CIGAR did not include H or S tags. Unique alignments are reported for reads that either have a single primary alignment or reads that have a secondary alignment (XA tag) but one alignment has a maximum mapping quality score of 60. Reported values are averaged over $n$=45 samples. Paired one-sided t-tests were conducted with $n-1$ degrees of freedom. Parameters marked with '*' indicate the null-hypothesis that ARS-UCD1.2 would perform better than the extended reference, while those without marks indicate the reverse. All tests rejected the null hypothesis.

| Parameter | Extended reference | ARS-UCD1.2 | Difference | Stdev | t-statistic & p-value |
|---|---:|---:|---:|---:|---|
| Unmap (%) * | 0.4291 | 0.4467 | -0.0176 | 0.00461087 | t =-24.12, p = 2.39e-25 |
| Alignment identity 99% (%) | 87.2716 | 87.1875 | 0.0841 | 0.00433417 | t = 122.72, p = 2.19e-52 |
| Alignment perfect (%) | 68.5732 | 68.4687 | 0.1045 | 0.00667272 | t = 99.04, p = 9.10e-49 |
| Clipped alignment (%) * | 2.1335 | 2.1923 | -0.0588 | 0.00891613 | t = -41.74, p = 2.81e-34 |
| Supplementary alignment (%) * | 0.2078 | 0.2219 | -0.0141 | 0.00379671 | t = -23.45, p = 6.69e-25 |
| Unique alignment (%) * | 83.2919 | 83.6016 | -0.3017 | 0.03348539 | t = -58.51, p = 6.50e-40 |

### Supplementary Note 1

#### Assembly of the Original Braunvieh (OBV) genome

The Original Braunvieh primary assembly was generated from PacBio HiFi CCS reads (study accession PRJEB42335 under sample accession SAMEA7759028), generated from subreads with minimum three passes and minimum predicted read quality of 20. The fastq data contained 86.9 gigabases, corresponding to nearly 30-fold coverage. The CCS reads were filtered by fastp [0.21.0] [3] with minimum average quality of Q20 and minimum read length of 1kb, with 99.99% of the data passing these thresholds. Hifiasm [v0.13-r308][4] was then used to generate the assembly from the reads using the additional parameters "-r 4 -a 5 -n 5" on a computing cluster. Hifiasm yields the primary contigs in the GFA format, which were then converted using gfatools [0.4-r196-dirty] into a fasta sequence representation. These contigs were then scaffolded using RagTag [v1.0.1][5] to the ARS-UCD1.2 reference, with custom parameters "--mm2-params "-c -x asm5" -r -m 1000000".

The contigs were validated for contiguity, completeness, and correctness by multiple independent tools, available in a Snakemake [5.26.1][6] pipeline online at https://github.com/AnimalGenomicsETH/bovine-assembly. Basic contiguity was determined through the *asmstat* command of paftools[7]. Similarly, the NGA50 value was determined through mapping the contigs to the ARS-UCD1.2 reference, and subsequently considering the length of alignment blocks again with *asmstat*. These values are described in Table SN1.

**Table SN1: Contiguity metrics of the primary Original Braunvieh assembly.**
Size refers to the total number of bases in the chromosomes and unplaced contigs. NG50 was calculated for both the contig set and the scaffolded assembly with the expected genome size taken from the ARS-UCD1.2 reference. Similarly, NGA50 is the NG50 value for aligned blocks of the assembly to the ARS-UCD1.2 reference.

| | Size | Contig NG50 | NGA50 | Scaffold NG50 | L50 | Contigs |
|---|---|---|---|---|---|---|
| **assembly** | 3.17gb | 86.0 | 68.9 | 96.3 | 15 | 765 |

Completeness of the assembly was determined through two independent approaches, BUSCO[8] and the *asmgene* command of paftools. The former relies on the OrthoDB datasets, specifically version 10 of the cetartiodactyla lineage. The latter uses cDNA libraries of annotated gene sequences from the ARS-UCD1.2 reference available from Ensembl. Both methods report a high completeness (>96%) with respect to predicted gene content, as shown in Table SN2.

**Table SN2: Predicted single-copy gene completeness of the primary Original Braunvieh assembly.**
Single-copy refers to genes that were correctly present once in assembly, while duplicates are genes which appeared more than expected. Fragmented genes are those which are only partially mapped, or fully mapped but split into multiple pieces. Missing genes are either not found or mapped below 10% of the expected gene.

| | Single copy | duplicates | fragmented | missing | total |
|---|---|---|---|---|---|
| **Busco** | 12533 | 283 | 166 | 353 | 13335 |
| **asmgene** | 18503 | 166 | 68 | 136 | 18873 |

Correctness was likewise determined by two k-mer based approaches, yak [r58][4] and Merqury[9]. Yak uses an approximate hash-table approach, while Merqury can be run in an exact mode. Both used short read sequences (2x150 bp) from the primary animal, which importantly were not used in generating the assembly, allowing for an independent evaluation. In addition, short read sequences from both parents enabled a quantification of the switch error rate. Only yak provided an estimate of the Hamming error rate, while only Merqury provides phased block statistics. An overview of these statistics is shown in Table SN3.

**Table SN3: K-mer based, reference-free validation of the primary Original Braunvieh assembly.**
Assembly quality value (QV) is given as a Phred quality score. Completeness estimates how many k-mers present in the short reads are found in the assembly contigs. The switch error rate is calculated differently by *yak* and Merqury, measuring the percent of wrongly phased adjacent SNPs in yak while in Merqury it measures the percent of wrongly phased haplotype-specific k-mers (“hap-mers”). There are more than 100 phase switches within a 20kb window (long-range switch). The Hamming error is the percent of SNP sites that are phased wrongly. The phase block statistic is the N50 after contigs have been broken at long-range switches, defined as more than 100 wrongly phased hap-mers per 20kb window.

| | **QV** | **Completeness** | **Switch error** | **Hamming error** | **Phased N50** |
|---|---|---|---|---|---|
| **Yak** | 48.76 | 100 | 0.012 | 0.37 | - |
| **Merqury** | 50.85 | 93.46 | 0.08 | - | 2.5 mb |

Furthermore, the assembly was validated by comparing structural variants called by pbsv [2.4.0] between the reads and the ARS-UCD1.2 reference and those called by mumandco [v2.4.2][10] between the assembly and the reference. There was good concordance between these approaches, for example an 8kb inversion identified in chromosome six of the assembly matched an 8kb inversion predicted by the read mapping.

The repeat content of the assembly was also in line with expectations, with approximately 48% of the assembly consisting of repeat elements or low complexity regions according to RepeatMasker version 4.1.1[11] using the Repbase repeat database (release 20181026)[12]. Several bovine-specific repeats were identified, along with telomeric or centromeric sequences not present in the existing ARS-UCD1.2 reference, indicating that several contigs are approaching chromosomal-scale and completeness.

#### Supplementary Note 2

##### Determination of the core and flexible parts of the pangenome graph

To investigate if the order of assemblies used to establish the multi-assembly graph impacts the core and flexible parts, we added the assemblies randomly to the graph. Core genome represents bases shared across all assemblies in the graph, while flexible genome represents number of bases that are variable across assemblies (i.e., not found in all assemblies)[13]. The pangenome increased gradually with the number of assemblies added, driven by an increase in the flexible genome. The core genome size decreased from 2480 Mb to 2400 Mb in the full graph, indicating that more genomic segments are variable across bovine species as we add more assemblies into the graph.

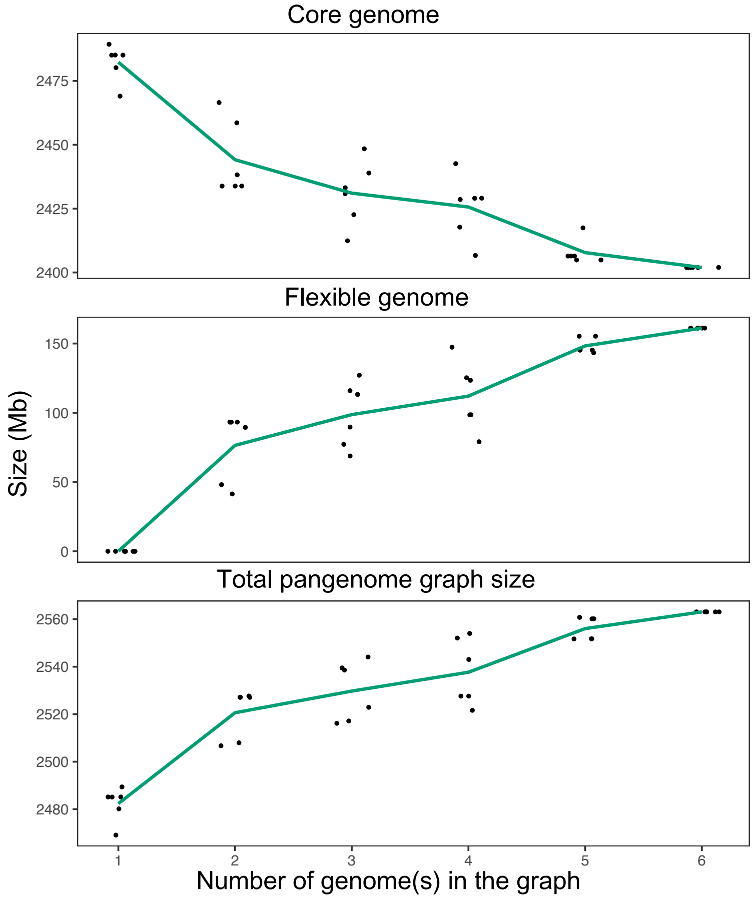


**Figure SN1. Profile of the multi-assembly graph with an increasing number of genomes integrated into the graph.** We varied the order and number of genomes added to the graphs, and calculated the number of bases in the pangenome, number of bases that are shared across all assemblies in the graph (core genome), and the number of bases that are variable across assemblies (i.e., not found in all assemblies, flexible genome). Points and lines indicate individual and average values.

Next, we investigated the profile of a multi-assembly graph that gradually increases in complexity. We built taurine-only graphs that contained either all or all but one taurine assemblies, a TauInd (four taurine and one indicine), and a full graph (four taurine, one indicine, and yak). The profile of the pangenome changed markedly as more distant assemblies were added to the graph. For example, the flexible part declined substantially from 6.10% in the full graph to 3.83% and 2.76% for the TauInd and taurine-only graph, respectively. However, when an individual taurine assembly is removed from the taurine-only graphs, the size of the flexible part changes only slightly (Figure SN2).

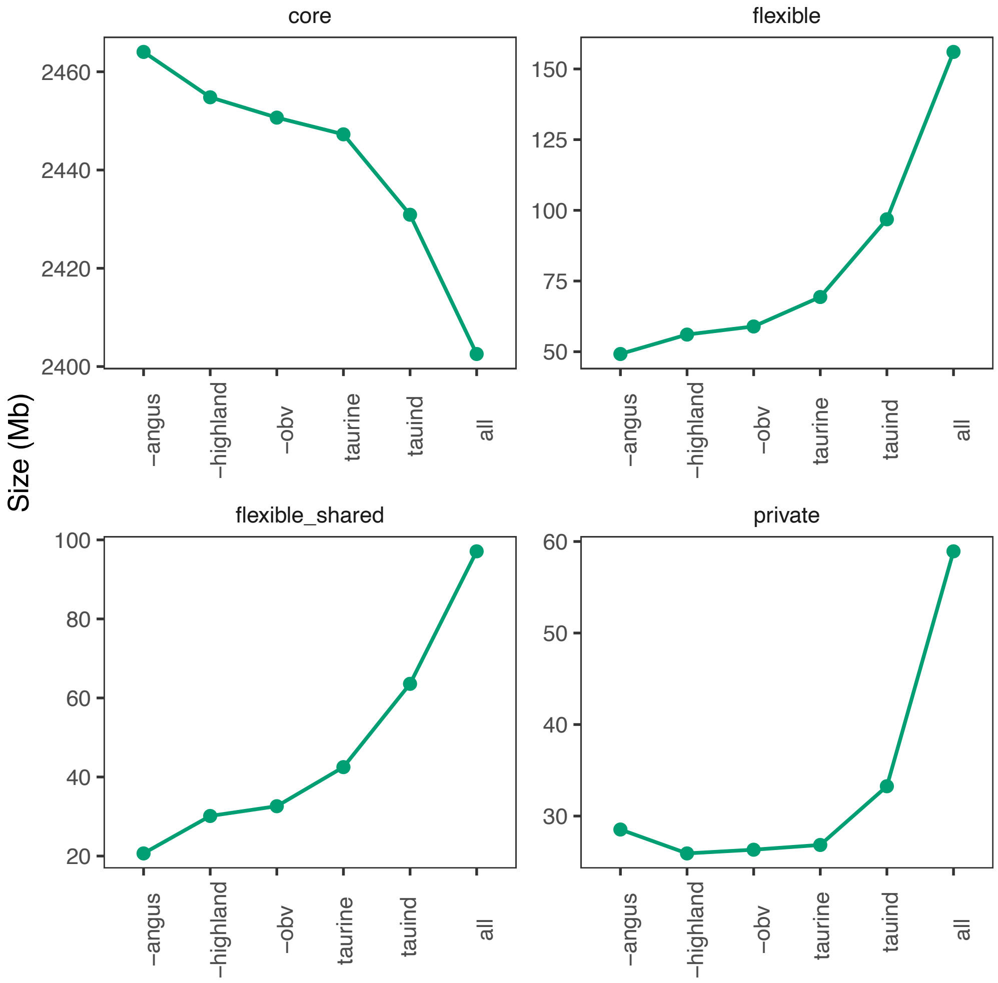


**Figure SN2. Pangenome profile as more distant assemblies are added to the graph**. The X-axis indicates the constructed graphs (- denotes the taurine assembly that was removed from the taurine-only graph, taurine denotes a graph with four taurine assemblies, the indicine graph contains all taurine assemblies and the assembly of Brahman, and all reflects a multi-assembly graph that contains all six assemblies (taurine, indicine, and yak). The core part is the size of the segments that are common to all assemblies, flexible_shared indicates the size of segments shared by at least two but not all assemblies, and private denotes the size of segments found only in a single assembly, thus flexible genome is composed of flexible_shared + private segment.

#### Supplementary Note 3

##### Differential expression analysis

We tested 13,085 genes that were expressed >= 1 CPM in at least eight samples (sample size from each group) for differential expression between *Mycobacterium bovis*-infected and non-infected control animals. We detected (adjusted FDR ≤0.05) 1,769 and 1,877 genes that were up-and down-regulated respectively in peripheral blood leukocytes of *Mycobacterium bovis*-infected cattle (Figure SN3). Of 12,813 genes of the Ensembl ARS-UCD1.2 genome annotation that were expressed at >=1 CPM in at least eight samples, 3610 (28.17%) were differentially expressed. Of 272 putatively novel genes that were expressed >=1 CPM in at least 8 samples, 36 (13.23%) were differentially expressed.

We found that genes relevant for the immune response were among the top differentially expressed genes with the greatest mean log-fold change (e.g., DEFB10 -8.24-fold, CXCL10 -3.30-fold, IL12B -3.11-fold, CXCL5 7.11-fold, CTLA4 4.25-fold, and CXCL8 5.70-fold), matching observations on an older reference genome annotation by McLoughlin *et al.*[14]. Multidimensional scaling (MDS) representations of transcript abundance estimates from either all 13,085 genes (Figure SN4b) or 3646 differentially expressed genes (Figure SN4a) separated *Mycobacterium bovis*-infected from healthy cattle. We discovered more differentially expressed genes of the Ensembl ARS-UCD1.2 genome annotation than McLoughlin *et al.* [14] (3610 vs. 3250), likely due to a vastly improved genome assembly (27,115 vs. 24,616 genes are included in build 101 (ARS-UCD1.2) and build 73 (UMD3.1), respectively).

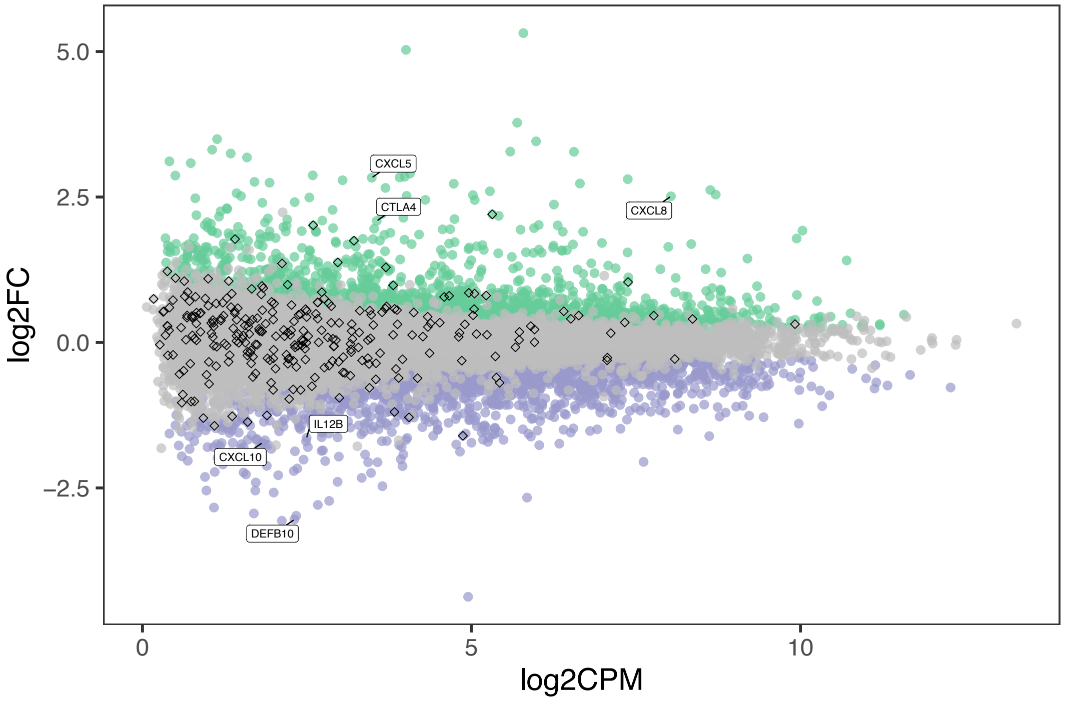


**Figure SN3. Smear plot from the differential expression analysis.** Grey, green, and purple color indicates genes with no expression difference, significant up-regulation, and down-regulation in peripheral blood leukocytes of *Mycobacterium bovis*-infected cattle. Diamonds indicate 272 putatively novel genes assembled from RNA sequencing reads mapping to non-reference sequences. Six genes reported by (McLoughlin *et al.* 2014) are indicated with the text labels.

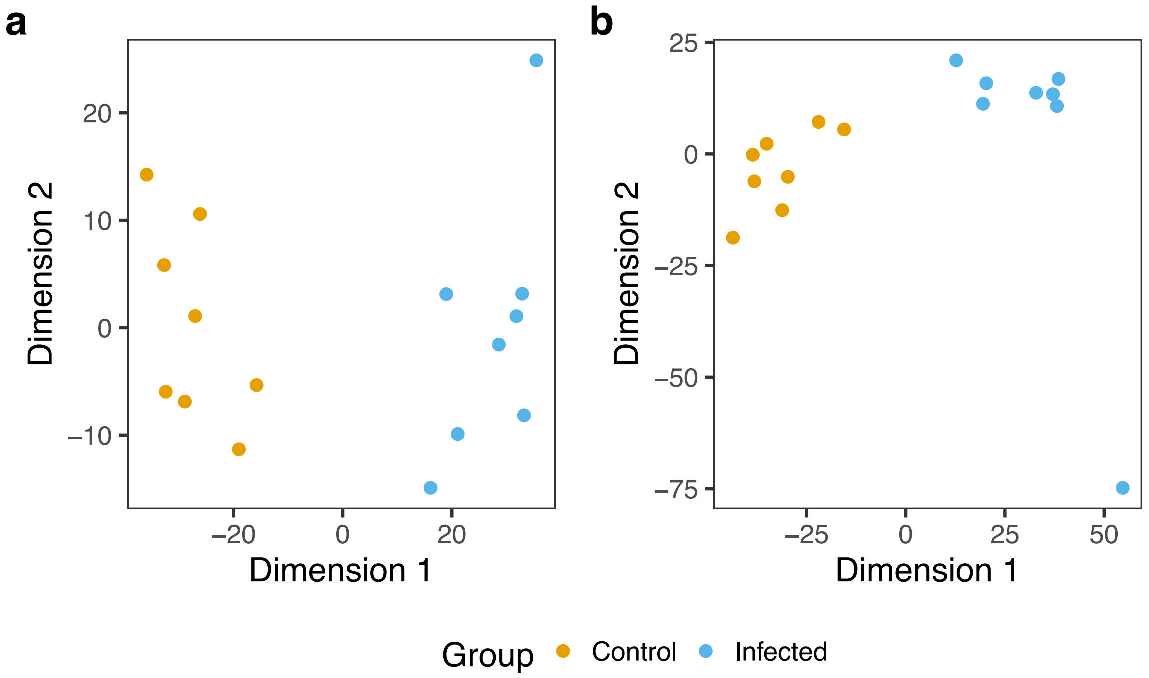


**Figure SN4. Multidimensional scaling analysis (MDS) based on transcript abundance estimates of (a) 3646 differentially expressed, and (b) 13,085 genes with CPM>=1 in eight samples.** Each point represents an individual *Mycobacterium bovis*-infected (blue) or control (orange) sample.
